# Podocalyxin and ciliary neurotrophic factor receptor are novel components of the surfaceome of chondrogenic cells

**DOI:** 10.1101/2025.05.19.654853

**Authors:** Patrik Kovács, Peter Brazda, Tibor Hajdú, Boglárka Harsányi, Krisztián Juhász, Roland Takács, Judit Vágó, Zhangzheng Wang, Clare Coveney, David J. Boocock, Csaba Matta

## Abstract

**Background:** Osteoarthritis is a degenerative joint disease characterized by progressive loss of articular cartilage and limited capacity for intrinsic repair. A major barrier to developing effective regenerative strategies is the incomplete understanding of the molecular mechanisms regulating chondrogenesis and cartilage maintenance. Cell surface proteins are key mediators of extracellular communication, adhesion, and signaling, yet the chondrogenic surfaceome remains incompletely mapped, with prior studies focusing primarily on mature or cytokine-activated chondrocytes. The aim of this study was to provide a temporal profile of the surfaceome during *in vitro* chondrogenic differentiation and to identify novel membrane proteins with potential roles in cartilage biology.

**Methods:** We applied a sialoglycoprotein-targeted glycocapture strategy to selectively enrich plasma membrane proteins from chick embryonic limb bud-derived micromass cultures undergoing chondrogenesis. Enriched samples underwent high-resolution shotgun proteomic analysis, and differentially expressed candidates were validated by western blotting, immunocytochemistry, and transient gene silencing. Functional effects on extracellular matrix gene regulation were assessed by quantitative RT-PCR and matrix histochemistry.

**Results:** This approach generated the first temporal surfaceome map of chondrogenic progenitors. Among identified candidates, two proteins not previously linked to chondrogenesis, podocalyxin (PODXL) and ciliary neurotrophic factor receptor (CNTFR), were detected at the plasma membrane and confirmed at the protein and transcript levels. Both proteins exhibited time-dependent downregulation during differentiation. Targeted knockdown revealed differential regulation of the fibrocartilage marker *COL1A1* expression, indicating non-redundant roles in cell-matrix signaling and survival pathways. Single-cell transcriptomic meta-analysis confirmed expression of both proteins in discrete human articular chondrocyte subpopulations.

**Conclusions:** This study expands the molecular framework of chondrogenesis, identifying PODXL and CNTFR as novel, temporally regulated surfaceome components with distinct roles in extracellular matrix signaling. These findings complement prior proteomic analyses of cytokine-activated mature articular chondrocytes and suggest new candidates for developmental cartilage biomarkers and therapeutic targets for osteoarthritis. Our results provide a resource for future cross-species surfaceome studies and highlight key pathways for further investigation into cartilage lineage specification and matrix adaptation.

**Plain English Summary:** Osteoarthritis is the most common joint disease worldwide, causing pain, stiffness, and reduced mobility as the smooth cartilage that covers the ends of bones gradually breaks down. Cartilage has a very limited ability to heal itself, and there are currently no treatments that can stop or reverse the disease. Scientists are working on new ways to repair or replace damaged cartilage, but to do this effectively, they need to understand the molecular “language” that cartilage-forming cells use to develop and maintain healthy tissue.

In this study, we examined the set of proteins that are present on the surface of cartilage-forming cells, known as the “surfaceome.” These surface proteins play important roles in how cells connect to their surroundings, receive signals, and work together to build the cartilage matrix. Using a method to selectively capture and analyze these proteins, we discovered two that had never been linked to cartilage development: podocalyxin and ciliary neurotrophic factor receptor.

We found that these two proteins may have opposite effects on cartilage quality. Because they can be detected on the outside of cells, they could be developed into markers for early osteoarthritis detection or as targets for new therapies. Our findings open the door to new strategies for diagnosing and treating joint disease by focusing on cell-surface communication.

## Background

Osteoarthritis (OA) is the most common form of arthritis, characterized by the progressive degeneration of articular cartilage, subchondral bone sclerosis, synovial inflammation, osteophyte formation, and subsequent loss of joint function, leading to chronic pain and disability (1). OA imposes a substantial global health burden, affecting more than 303 million adults worldwide (2). Despite its high prevalence, there are currently no FDA-approved disease-modifying OA drugs capable of halting disease progression or promoting cartilage repair, largely due to the heterogeneous nature of OA and incomplete understanding of its molecular drivers (3). Current treatments remain palliative, focusing on symptom relief, underscoring an urgent unmet need for innovative therapeutic strategies (4).

Regenerative medicine has emerged as a promising avenue to address this gap. Stem cell-based approaches, particularly those using mesenchymal stem cells (MSCs), hold potential for cartilage regeneration and functional restoration (5). However, clinical outcomes have thus far been modest (6). The major obstacles include poor MSC survival in the inflammatory joint microenvironment, coupled with limited integration into native cartilage, which often result in transient improvements without sustainable extracellular matrix (ECM) restoration (2), (7). A deeper understanding of the molecular pathways that control chondrogenesis is therefore essential for improving cartilage repair outcomes (8).

A key step in this process is identifying novel biomarkers in chondroprogenitor cells that could provide mechanistic insight into cartilage differentiation, guide patient stratification in light of OA heterogeneity, and inform targeted regenerative strategies. Biomarkers predictive of chondrogenic capacity could significantly improve MSC-based cartilage repair by enabling optimal cell sourcing and enhancing survival, integration, and ECM restoration.

In this context, profiling the surfaceome, i.e., the full dynamic complement of cell surface proteins, is of particular importance for understanding chondroprogenitor cell biology and optimizing cartilage tissue engineering approaches (9-11). Current surface markers such as CD44, CD73 show variable association with the chondrogenic potential of MSCs derived from different tissues (12), highlighting the need for more reliable and functionally relevant surface markers (13). Such biomarkers could not only improve the selection of optimal cell types for therapy but also support the design of biomaterials that mimic native chondrogenic niches (14).

Our group has previously performed sialoglycoprotein-based surfaceome mapping of mesenchymal stem cells and migratory chondrogenic progenitor cells to define adhesive and signaling signatures at early cartilage lineage commitment, and to highlight differences between these two cell types (9). In a subsequent study, we used a similar approach on primary equine articular chondrocytes under normal and pro-inflammatory conditions to reveal cytokine-induced surfaceome remodeling (15). While such earlier work has defined features of the chondrocyte surfaceome under various physiological and pathological conditions, there is a need for systematic temporal analysis during chondrogenic differentiation. Therefore, by building on earlier findings, we aimed to establish a temporal, quantitative map of the chondrogenic surfaceome using a developmental *in vitro* model to identify new membrane-associated proteins involved in cartilage differentiation and phenotype stabilization.

Here, we present a temporal landscape of the surfaceome of chondrogenic progenitor cells during chondrogenesis. Using a sialoglycoprotein-targeted glycocapture method with aminooxy-biotin, followed by shotgun proteomics (9, 15, 16), we enriched the cell surface subproteome. We aimed to determine which membrane-associated proteins may be associated with the transition from chondroprogenitor to mature chondrocyte states and how these proteins integrate with extracellular matrix and cell-survival signaling. We identified proteins not previously linked to chondrogenic differentiation. Notably, we discovered podocalyxin (PODXL), a sialomucin known for regulating podocyte polarity, and ciliary neurotrophic factor receptor (CNTFR), a neurotrophic signal transducer, as novel constituents of the chondrogenic surfaceome. Their detection suggests convergence of adhesion- and survival-related signaling pathways during chondrogenesis, expanding current knowledge of extracellular signaling in cartilage development and highlighting potential translational targets for OA therapy.

## Methods

### 1. Experimental setup

This study employed a staged approach to define the surfaceome of embryonic chicken limb bud-derived chondrogenic cells at key stages of chondrogenic differentiation. Temporal proteomic profiling was conducted at days 1, 3, 6, 10, and 15 of micromass culture corresponding to key histodifferentiation stages (17, 18), combining sialoglycoprotein enrichment via aminooxy-biotin conjugation with high-resolution LC-MS/MS on 3 biological replicates (N = 3). Candidate surface markers ciliary neurotrophic factor receptor (CNTFR) and podocalyxin (PODXL) were validated through RT-qPCR, western blotting, and immunocytochemical localization. The integrated workflow, spanning discovery to functional validation, is summarized in Figure 1.

**Figure 1.**
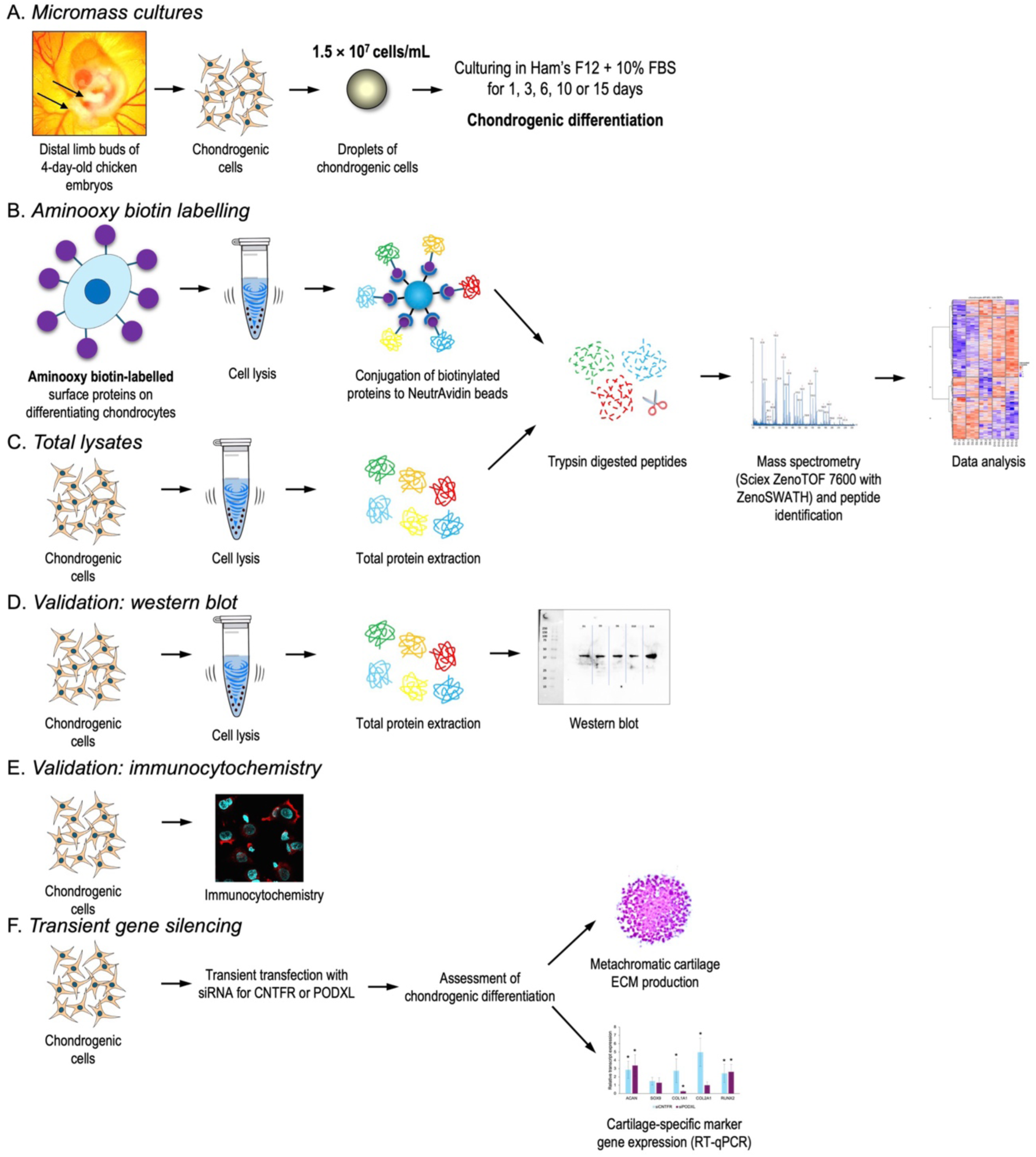
Experimental workflow.

### 2. Chondrogenic micromass cultures

Chondrogenic micromass cultures were established from embryonic chicken limb bud chondroprogenitors using a standardized protocol optimized for spontaneous chondrogenesis (18). Briefly, distal parts of limb buds were carefully excised from early-stage (Hamburger– Hamilton 22–24) chicken embryos and dissociated into a single-cell suspension via trypsin-EDTA digestion. Cells were resuspended in Ham’s F12 medium (Gibco, Thermo Fisher Scientific) supplemented with 10% fetal bovine serum (FBS, Gibco) and plated as 100 µL micromass droplets (1.5 × 10⁷ cells/mL) onto 6-well plates. Following a 2-hour adhesion phase (37°C, 5% CO₂), cultures were gently flooded with fresh medium to maintain 3D architecture. This model recapitulates *in vivo* chondrogenesis, including *COL2A1* upregulation and *SOX9*-dependent ECM synthesis, as previously characterized in this system (17, 18). Cultures were harvested at days 1, 3, 6, 10, and 15 to capture key transitions in surfaceome dynamics during early differentiation.

### 3. Cell surface proteome labelling and enrichment, and total cell lysates

For a detailed methodology for cell surface sialoglycoprotein labelling and enrichment of chondroprogenitor cells in micromass cultures, please see our protocol paper (16). Unless otherwise stated, all reagents were purchased from Thermo Fisher Scientific. Briefly, sialic acid residues on the cell surface proteins of micromass cultures were oxidized using 1 mM sodium meta-periodate to generate reactive aldehyde groups. This was followed by incubation in the dark for 1 hour at 4°C with 100 mM aminooxy-biotin (Biotium). After quenching with glycerol, the labelled cells were lysed (1% Triton X-100, 150 mM NaCl, 5 mM iodoacetamide, 10 µL protease inhibitor cocktail, 0.1 mg/mL PMSF and 10 mM Tris-HCl; pH 7.6). Biotinylated proteins were affinity-purified using NeutrAvidin agarose beads, followed by stringent washes to remove nonspecific binders. Captured protein were reduced by 100 mM DTT and alkylated in 50 mM iodoacetamide, and further wash steps were performed to minimize contamination from intracellular components, and to prepare the sample for LC-MS/MS analysis. Captured proteins were digested overnight with 5 μg MS-grade trypsin in 50 mM NH_4_HCO_3_. Solvents were then evaporated in a vacuum concentrator centrifuge and stored at –80°C.

Parallel total proteome analysis of micromass cultures were prepared using lysis in RIPA and sonication to shear DNA (3 × 10 sec pulses), ensuring membrane protein solubilization. Samples were then stored at –80°C. This approach enabled comparative quantification of surface-enriched vs. intracellular proteins during chondrogenesis.

### 4. Mass spectrometry, protein identification and quantification

Dried peptide samples were reconstituted in 150 μL of 10% acetonitrile with 0.1% formic acid, shaken at 5°C for 20 minutes, followed by addition of 150 μL 0.1% formic acid to yield a final concentration of 5% acetonitrile and 0.1% formic acid. After centrifugation at 12,000 × g for 2 minutes to remove particulates, supernatants were transferred to LC-MS compatible vials. Peptide separation was performed on a Waters M-Class UPLC system equipped with a Phenomenex Kinetex XB-C18 column (2.6 μm, 15 × 0.3 mm) maintained at 30°C, using a linear gradient of 3–35% acetonitrile (mobile phase B) over 12 minutes at a flow rate of 10 μL/min, followed by washing at 80% B and re-equilibration to 3% B. Three microlitres of each sample were injected in direct inject mode. Mass spectrometric analysis was conducted on a Sciex 7600 ZenoTOF operating in positive ion mode with Data Independent Acquisition (DIA, zenoSWATH), employing a 25 ms TOF-MS scan and 65 variable SWATH windows (12 ms each) covering m/z 400–750, with a total cycle time of 1.146 s. Raw data files were processed using DIA-NN (version 1.8.1) (19), with a *Gallus gallus* UniProt FASTA database (Oct 2021), enabling library-free search and deep learning-based spectra prediction. Label-free quantification was performed at a precursor false discovery rate (FDR) of 1.0%, with default DIA-NN parameters unless otherwise specified. Protein identification and quantification results were exported, and proteins were annotated using Gene Ontology (GO) information from UniProt to infer subcellular localization (16).

The mass spectrometry proteomics data have been deposited to the ProteomeXchange Consortium *via* the PRIDE partner repository with the dataset identifier PXD064013.

### 5. Bioinformatics and pathway analysis

Proteomic intensity data were log2-transformed. Missing values were imputed using a left-censored normal distribution via the minProb method (q = 0.001) from the *promor* package in R (20) to simulate low-abundance signals below the detection threshold. Entries representing protein families (where individual proteins could not be reliably deconvoluted) were excluded from the downstream bioinformatic analysis workflow, as some of these were also present as single entries in the matrix. Principal component analysis (PCA) was used to assess sample variability and potential batch effects.

Differential protein expression (DEP) analysis was performed using the *DEqMS* package in R, applying linear modeling with variance moderation based on peptide counts. Proteins with |log2 fold-change| > 1 and Benjamini-Hochberg adjusted p-value (FDR) < 0.05 were considered DEPs. The *gorth* package was used to perform ortholog mapping from chicken (*Gallus gallus*) to human (*Homo sapiens*). The resulting set of DEPs was z-score scaled and grouped using hierarchical clustering to identify temporal expression patterns.

Gene Ontology (GO) enrichment analysis was performed for each DEP list and temporal DEP cluster using *enrichGO*() and *compareCluster*() (*org.Hs.eg.db*, p-value < 0.05, q-value < 0.2), covering Biological Process (BP), Molecular Function (MF), and Cellular Component (CC) categories. Statistical analysis and visualization were carried out in R.

Our NGS dataset (BioProject IDs: PRJNA817177; PRJNA938813) (17) was processed using R. Weighted gene correlation network analysis (WGCNA) was performed with the *WGCNA* R package. For the construction of the protein-protein interaction (PPI) network, we utilized the STRING database, and Cytoscape was employed to further screen the proximal associated proteins. Enrichment analysis was carried out using the *clusterProfiler* R package.

### 7. Immunocytochemistry

On specified culture days, cells were dissociated using mild collagenase type I treatment (Sigma) for 30 minutes at 37°C, resuspended in fresh F12 medium, and seeded onto glass coverslips in 24-well plates. Cells were allowed to adhere for 2 hours in a humidified incubator (37°C, 5% CO_2_). Following attachment, cells were fixed in 10% formalin in 1 × PBS for 30 minutes at room temperature (RT). After washing in PBS, non-specific binding sites were blocked with 3% bovine serum albumin (BSA) and 10% serum from the host species of the secondary antibody (see below) in PBS for 1 hour at RT.

For immunostaining, cells were incubated overnight at 4°C with anti-CNTFR (Abcam; Cat. No. ab127425) at a dilution of 1:500, and anti-PODXL (Proteintech; Cat. No. 18150-1-AP) antibodies at a dilution of 1:500, diluted in blocking solution. Primary antibodies were detected using a goat anti-rabbit IgG secondary antibody conjugated to Alexa Fluor 555 (Life Technologies Corporation; Cat. No. A27039) at a dilution of 1:1000, incubated for 1 hour at RT. Cultures were mounted in Vectashield Hard Set mounting medium (Vector Laboratories) containing DAPI to visualize nuclei. Subcellular localization of PODXL and CNTFR was analyzed using an Olympus FV1000S confocal microscope (Olympus) equipped with a 60× oil immersion objective (NA: 1.3). For excitation of AlexaFluor 555, 543 nm laser line was used. The average pixel dwell time was 4 µs. Images of Alexa Fluor 555 and DAPI were overlaid.

### 8. Western blot analysis

Total cell lysates for SDS–PAGE were prepared as previously described (21). To ensure equal protein loading, after protein concentrations were determined, four-fold concentrated Laemmli buffer (Thermo Fisher Scientific) was added to the lysates, and samples were heated at 95°C for 10 minutes. In each lane, 20 μg of protein per sample was loaded onto 10% SDS–polyacrylamide gels for separation. Following electrophoresis, proteins were transferred onto nitrocellulose membranes using electroblotting. Membranes were blocked with 5% non-fat dry milk dissolved in PBS for 1 hour at RT. Subsequently, membranes incubated overnight at 4°C with the primary antibodies (anti-CNTFR at a dilution of 1:500, and anti-PODXL at a dilution of 1:500). After three washes in PBST (PBS containing 0.1% Tween-20) for a total of 30 minutes, membranes were incubated with an anti-rabbit IgG secondary antibody (Bio-Rad Laboratories) in 1:1000 dilution for 1 hour at RT. Protein bands were visualized using enhanced chemiluminescence (Millipore) according to the instructions provided by the manufacturer. Equal protein loading was verified by probing for β-actin as a loading control.

### 9. Transient gene silencing of CNTFR and PODXL

To investigate the impact of transient gene silencing of *CNTFR* and *PODXL* on chondrogenic differentiation, freshly isolated limb bud-derived progenitor cells were transfected with siRNA. Cells were resuspended at a density of 1.5 × 10⁷ cells/mL in Ham’s F12 medium and seeded into 12-well plates. Custom-designed siRNAs targeting CNTFR (Silencer™ Custom Designed siRNA; Cat. No. 4399666, Design ID: ABD1VPK) and PODXL (Design ID: ABHSP78), along with a non-targeting (NT) control (Silencer™ Select Negative Control No. 1 siRNA; Cat. No. 4390844; Life Technologies), were transfected into chondrogenic cells 2 hours post-attachment using Lipofectamine™ RNAiMAX Transfection Reagent (Cat. No. 13778150; Life Technologies) and Opti-MEM™ Reduced Serum Medium (Cat. No. 51985034; Life Technologies).

For the analysis of cartilage ECM production following gene silencing, colonies were fixed on culture day 6, stained with DMMB (pH 1.8), and photographed. Cartilage ECM production was quantified using a MATLAB script (18), with values normalized to NT control levels. Relative cell numbers or viability (inferred from mitochondrial activity) were analyzed via MTT assay (18). Optical density values were expressed as percentage changes relative to NT control.

Gene expression profiling of chondrogenic marker genes was performed as described previously (17) with minor adjustments. Micromass cultures were harvested 48 hours post-transfection. As cartilage ECM accumulation generally requires several days beyond this initial time point, these assays were designed to capture transcriptional trends reflecting early modulatory effects on chondrogenic gene regulation rather than fully developed phenotype changes. Total RNA was isolated using TRI Reagent (Applied Biosystems) following to the manufacturer’s instructions. Total RNA (1 μg) was reverse-transcribed into cDNA using the High-Capacity cDNA Reverse Transcription Kit (Thermo Fisher Scientific) according to the manufacturer’s protocol. The expression patterns of chondrogenic marker genes were quantified via CYBR Green-based RT-qPCR using the 2^−ΔΔCt^ method. Primer sequences (Integrated DNA Technologies) are provided in Table S1 (Additional file 1). RT-qPCR reactions were set up using GoTaq qPCR Master Mix (Promega) and 20 ng of input cDNA per each 10-μL reaction. Reactions were run on a QuantStudio 3 System (Thermo Fisher Scientific) using a standard thermal profile (17). Of the six reference genes evaluated for stability (*HPRT1*, *PPIA*, *RPL4*, *RPL13*, *RPS7*, *YWHAZ*), BestKeeper (22) identified *RPS7* as the most stable transcript. All target gene data were normalized to *RPS7* expression.

## Results

### 1. Temporal proteome dynamics in chondrogenic cultures

Principal component analysis (PCA) performed on the 5,207 proteins/protein groups identified in total lysates from chondrogenic micromass cultures revealed distinct proteomic profiles at each differentiation stage, with biological replicates clustering tightly by culture day (Figure 2A–B). The first principal component (PC1), which explains 42% of the variance, captured temporal progression, separating undifferentiated (day 1) from differentiated (days 10 and 15) cultures. PC2 (19% variance) distinguished intermediate differentiation stages (days 3–6) from both early and late time points.

**Figure 2.**
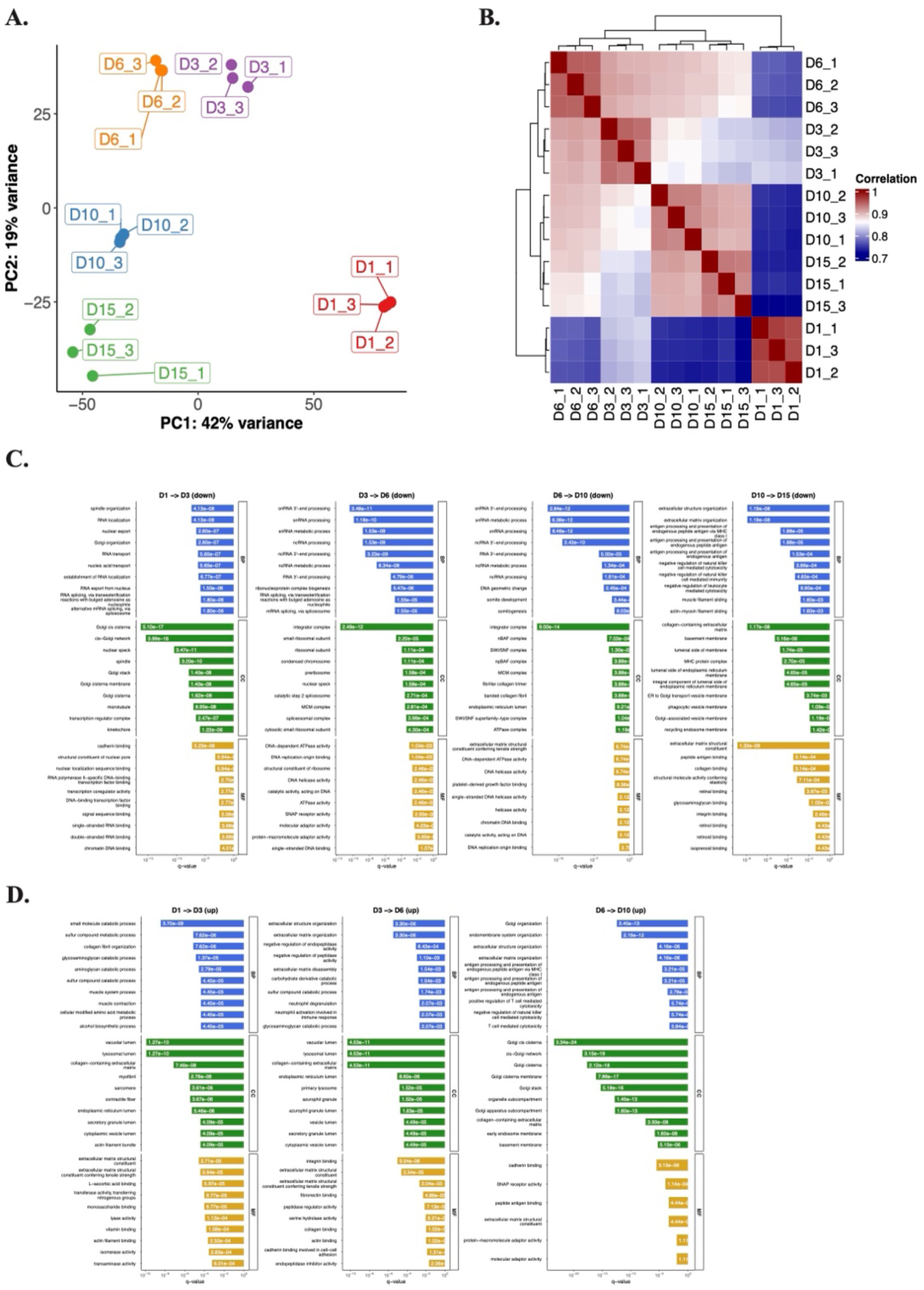
Quantitative proteomics analysis of total cell lysates from chondrogenic micromass cultures. **A.** Principal component analysis (PCA) of the total lysate samples of micromass cultures of chondroprogenitor cells undergoing chondrogenesis (5,207 proteins in the entire dataset). Each dot represents a biological replicate (n = 3). The PCA plot shows unsupervised clustering according to culture age along the principal component (PC) 1 and PC2 hyperplanes, accounting for 42% and 19% of the variance, respectively. **B.** The correlation matrix also demonstrates clear separation among the samples according to culture age. **C–D.** Enrichment analysis using gene ontology (GO) annotations for pairwise comparisons of culture days, showing the top GO terms (biological process (BP), cellular compartment (CC), or molecular function (MF)) by ranking them by the adjusted p-values (p.adjust < 0.001). Plots shown separately for upregulated (**C**, top panel) and downregulated (**D**, bottom panel) proteins. The *x*-axis is the adjusted p-value of the enriched terms.

A total of 792 proteins (323, 159, 272, and 38 for each consecutive timepoint comparison: day 1–3, day 3–6, day 6–10, and day 10–15, respectively) were identified as significantly upregulated. Similarly, 1368 proteins (714, 231, 178, and 245 for each corresponding timepoint comparison) were identified as significantly downregulated (Figure S1 in Additional file 1). Full lists of differentially expressed proteins (DEPs) for pairwise comparisons between consecutive days and between undifferentiated (day 1) versus mature (day 15) cultures are provided in the Supporting Information (Additional file 2).

To better characterize DEPs between each time point, we performed gene ontology (GO) analysis on the DEPs (Figure 2C–D). In case of proteins that were upregulated in consecutive time points, GO terms related to collagen-containing ECM synthesis, ECM organization, and integrin binding were over-represented. In contrast, when we looked at the downregulated DEPs separately, a much broader range of GO terms were identified, such as those related to DNA and RNA metabolism, DNA replication, and transcription factor binding, especially in the earlier time points. Between days 10–15, ECM-related GO terms were also enriched, indicating that ECM production has been completed by day 15 (mature/hypertrophic) cultures.

### 2. Representative proteins and pathways in chondrogenic cultures

The identified DEPs in the total lysates of chondrogenic cultures were categorized into 5 clusters to delineate expression dynamics (Figure 3A–B). Protein lists for each cluster are provided in the Supporting Information (Additional file 3).

**Figure 3.**
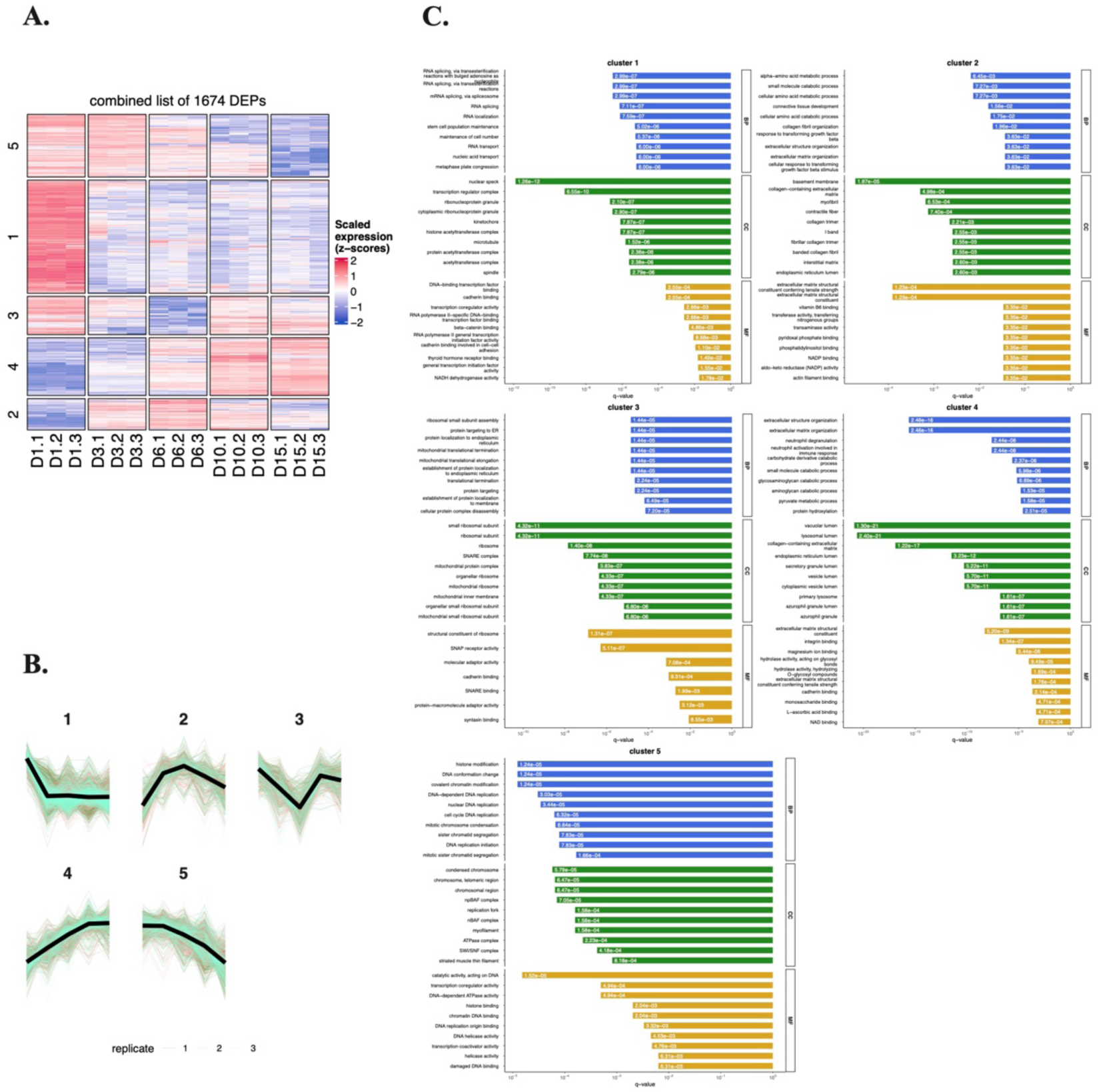
Unsupervised clustering analysis of differentially expressed proteins (DEPs) in total cell lysates of chondrogenic cultures using the k-means algorithm. A–B. The expression dynamics of each cluster are shown in clusters 1–5. *x*-axis, time points (days of culturing). **C.** Pathway analysis using Gene Ontology (GO) annotation for the 5 clusters of proteins in total lysates of chondrogenic cells undergoing chondrogenic differentiation. Select terms for GO biological process (BP), cellular component (CC), and molecular functions (MF). Barplot shows significant terms, the *x*-axis is gene (protein) ratio and the number of proteins belonging to given enriched terms. These clusters outline a progressive remodeling of the proteome from a proliferative, transcriptionally active progenitor profile toward an ECM-producing, metabolically adapted chondrocyte phenotype.

**Cluster 1** (625 proteins) exhibited an initial steep downregulation during early *in vitro* chondrogenesis (days 1–3), plateauing thereafter. This group showed enrichment for GO terms related to RNA splicing, histone acetyltransferase activity, and RNA polymerase II mediated transcription (Figure 3C), suggesting phased suppression of transcriptional machinery in maturing cultures. Key proteins in this group include AKT1, FN1, SOX5, NOTCH1, and ITGB1. AKT1 is a central component in the PI3K/Akt pathway, which modulates the balance between chondrocyte proliferation and differentiation (23). Fibronectin (FN1), an ECM glycoprotein, drives chondrocyte differentiation and collagen production via the PI3K/Akt signaling pathway (24). The SOX5 transcription is part of the SOX trio (SOX5, SOX6, SOX9) driving chondrogenic lineage commitment and promoting expression of key cartilage ECM genes such as *COL2A1* (25). NOTCH1 is a receptor that regulates early chondrogenic lineage differentiation via SOX9 (26). Integrin beta-1 (ITGB1) is a cell surface receptor that mediates cell–matrix adhesion and signals for chondrocyte proliferation, migration, and differentiation (27).

**Cluster 2** (173 proteins) displayed peak expression at day 6, with overrepresented GO terms linked to amino acid metabolism, connective tissue development, and collagen synthesis. This cluster included several functionally relevant proteins outside the collagen family (such as COL1A1, COL5A1, COL12A1, COL14A1, COL15A1), notably FRZB, TGFBR2, NFATC1, and THBS2, all of which are recognized as being involved in chondrocyte phenotype regulation, matrix remodeling, growth factor signaling, and cartilage regeneration. Specifically, frizzle-related protein (FRZB) is a secreted Wnt agonist that maintains chondrocyte phenotype and is highly expressed in articular chondrocytes (28). Transforming growth factor beta receptor II (TGFRB2), a cell surface receptor, is a critical regulator of chondrocyte maturation, and is essential in maintaining cartilage integrity (29). Nuclear factor of activated T-cells 1 (NFATC1) is a transcription factor that acts as a marker in articular cartilage progenitor cells. It is a key mediator of chondrocyte fate (30). Thrombospondin 2 (THBS2) is an ECM glycoprotein involved in cartilage regeneration (31).

**Cluster 3** (211 proteins) followed a U-shaped trajectory: downregulated initially but upregulated in later stages and was enriched for GO terms related to mitochondrial and ribosomal functions. This cluster contained key signaling and matrix-interacting proteins, including ACVR1 (BMP receptor required for chondrogenesis), decorin (DCN), syndecan-1 (SDC1), tenascin-C (TNC), and transferrin receptor (TFRC), all of which have mechanistic roles in regulating chondrocyte proliferation, differentiation, matrix integrity, and developmental signaling in cartilage tissue (32, 33). Notably, decorin is a proteoglycan in the cartilage pericellular and territorial matrix that regulates ECM architecture and load-bearing properties (34). Syndecan-1 is a cell-surface protepglycan involced in cell–matrix interactions (35). Tenascin-C is an ECM glycoprotein with important roles in cartilage development and ECM production (36).

**Cluster 4** (319 proteins) demonstrated progressive upregulation, peaking in mature chondrocytes (days 10–15), suggesting that these proteins are associated with the later stages of chondrogenic differentiation and may contribute to the functional maturation of chondrocytes. Over-represented GO terms in this cluster were related to ECM and collagen synthesis. This cluster included classic and emerging modulators of cartilage ECM and chondrocyte function such as ACAN (aggrecan, the major cartilage proteoglycan), major collagen types (COL2A1, COL6A1, COL8A1, COL9A1, COL22A1), COMP (cartilage oligomeric matrix protein; involved in matrix assembly), MATN1 (matrilin-1; involved in mechanotransduction), TIMP3 (tissue inhibitor of metalloproteinases; involved in matrix protection), ENO1 (alpha-enolase; involved in glycolysis) and LDHA (lactate dehydrogenase; involved in chondrocyte-specific glycolytic adaptation), all vital for cartilage tissue integrity and repair (8, 37).

**Cluster 5** (346 proteins) showed a similar expression pattern to cluster 1 but with a more sustained downregulation throughout chondrogenesis. Over-represented GO terms in this cluster included histone modification and cell proliferation (DNA replication and chromosome condensation). This cluster included functional components and matrix regulators such as CD44 (hyaluronan adhesion), PDGFRA (platelet-derived growth factor receptor alpha; essential for chondroprogenitor specification), FBN2 (fibrillin-2; involved in matrix assembly and growth factor regulation), and VCAN (versican; modulates cartilage morphogenesis and growth factor localization), all implicated in cartilage and joint development, repair, or disease (38-41).

Next, we carried out a detailed protein-protein interaction (PPI) analysis of each cluster using STRING and generated a small network with the top 20 hub proteins (Figure S2). In the PPI network of **Cluster 1**, spliceosome and ribosome biogenesis are recurrent pathways. Components of the PI3K/Akt and Wnt signaling were also enriched, whereas BIRC5 and SUMO1 regulate apoptosis and post-translational modifications, respectively. The PPI network in **Cluster 2** is centrally enriched in ECM dynamics and cell–matrix signaling, with focal adhesion and PI3K/Akt as hub pathways. The proteins in the network of **Cluster 3** are predominantly associated with ribosomal biogenesis, translation, and RNA processing, with strong enrichment in conserved pathways such as global protein synthesis, protein processing in endoplasmic reticulum, and mitochondrial translation. In a similar way to Cluster 2, the PPI network of **Cluster 4** is enriched in ECM remodeling, glycolysis, and cell adhesion signaling. ECM dynamics dominate this cluster, with collagens and modifiers driving ECM integrity. Glycolysis enzymes highlight metabolic reprogramming, and focal adhesion and PI3K/Akt couple signaling to pro-survival pathways. For **Cluster 5**, the PPI network is enriched in cell cycle regulation, DNA replication/repair, and chromatin remodeling.

### 3. Chondrogenic surfaceome dynamics via aminooxy-biotin enrichment

We next profiled the surfaceome during chondrogenic differentiation. Of 522 quantified proteins/protein groups, 18 detected in only one time point were excluded, leaving 504 proteins for differential protein expression analysis. PCA revealed low degree of variability among biological replicates, with clear separation by culture day along PC1 (40% variance) (Figure 4). This trajectory mirrored trends observed in the total proteome, reflecting stage-specific surfaceome dynamics as cells progressed from undifferentiated (day 1) to mature (day 15) states.

**Figure 4.**
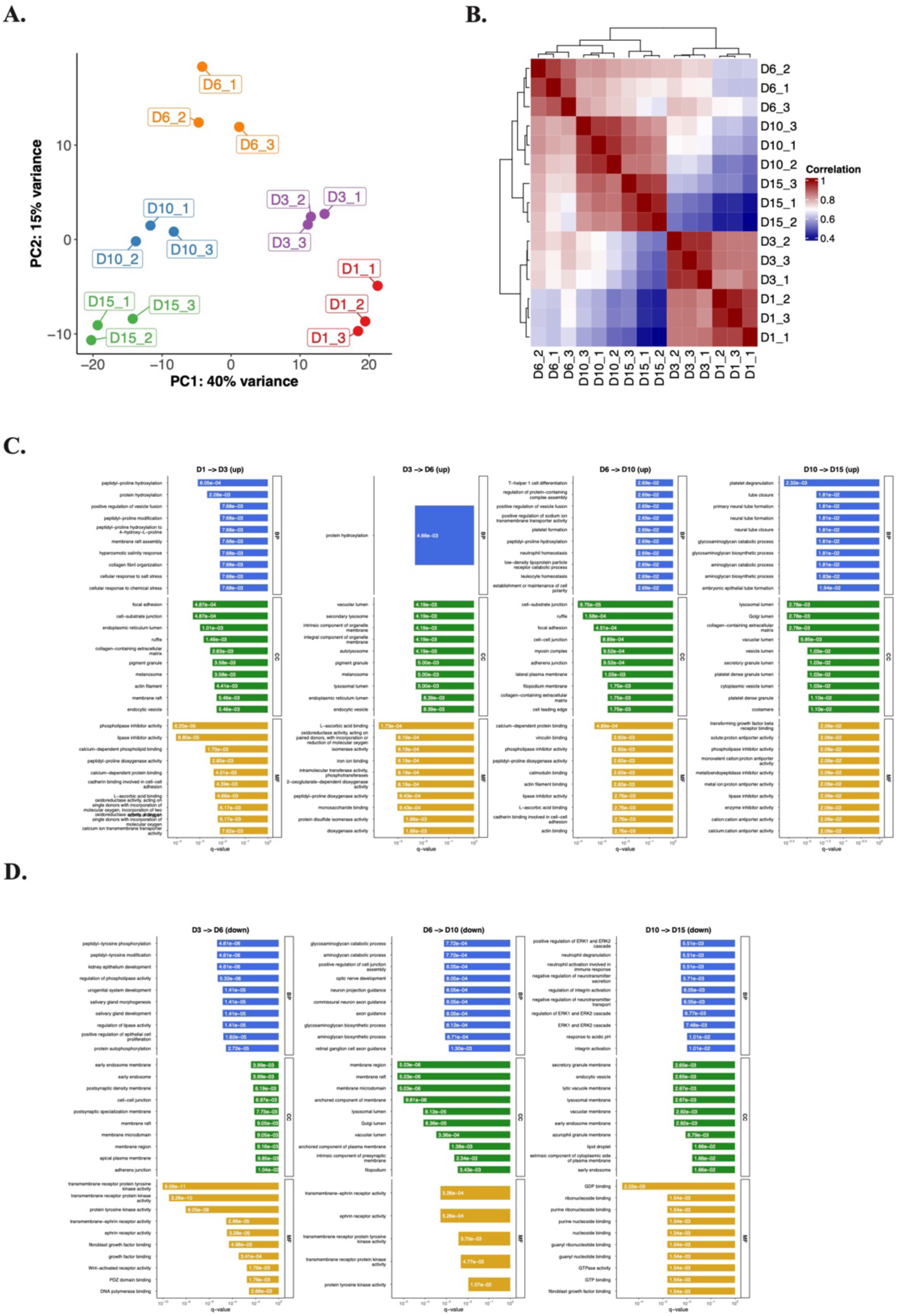
Quantitative proteomics analysis of aminooxy biotin (AOB) enriched samples from chondrogenic micromass cultures. **A.** PCA of the AOB-enriched samples of micromass cultures undergoing chondrogenesis (504 proteins in the dataset). Each dot represents a biological replicate (n = 3). The samples clustered together according to culture age along the principal component (PC) 1 and PC2 hyperplanes, which account for 40% and 15% of the variance, respectively. **B.** The correlation matrix (right panel) also signifies separation among the samples according to differentiation status. **C–D.** Pathway analysis using Gene Ontology (GO) annotation for the AOB-enriched subproteome of chondrogenic cells undergoing chondrogenic differentiation. Panel shows the top GO terms based on biological process (BP), cellular compartment (CC), or molecular function (MF), separately for upregulated (**C**, top panel) and downregulated (**D**, bottom panel) proteins, following ranking by adjusted p-values. The *x*-axis is the adjusted p-value of the enriched terms.

A total of 254 proteins (72, 74, 89, and 19 for each consecutive timepoint comparison: day 1–3, day 3–6, day 6–10, and day 10–15, respectively) were identified as significantly upregulated. Similarly, 170 proteins (10, 82, 31, and 47 for each corresponding timepoint comparison) were identified as significantly downregulated (Figure S3). DEP lists for pairwise comparisons of subsequent culture days are provided in the Supporting Information (Additional file 4).

We performed GO analysis on DEPs that were up or downregulated in later time points to better characterize DEPs between each time point (Figure 4C–D). Upregulated proteins were enriched in collagen biosynthesis (terms such as peptidyl-prolyl hydroxylation, L-ascorbic acid binding), ECM maturation (glycosaminoglycan biosynthesis), and cell–matrix adhesion, consistent with chondrocyte functional specialization. In contrast, GO terms associated with downregulated DEPs included developmental signaling (ephrin receptor pathway, Wnt-activated receptor activity, regulation of ERK1 and ERK2 cascade), growth factor modulation, and several terms related to neural processes (synapses and axonal growth), suggesting attenuation of plasticity pathways in mature cells.

### 4. Representative proteins and pathways in the chondrogenic surfaceome

Proteins in the aminooxy-biotin-enriched subproteome were classified into four clusters based on expression dynamics (Figure 5A–B). Protein lists for each cluster are provided in the Supporting Information (Additional file 5).

**Figure 5.**
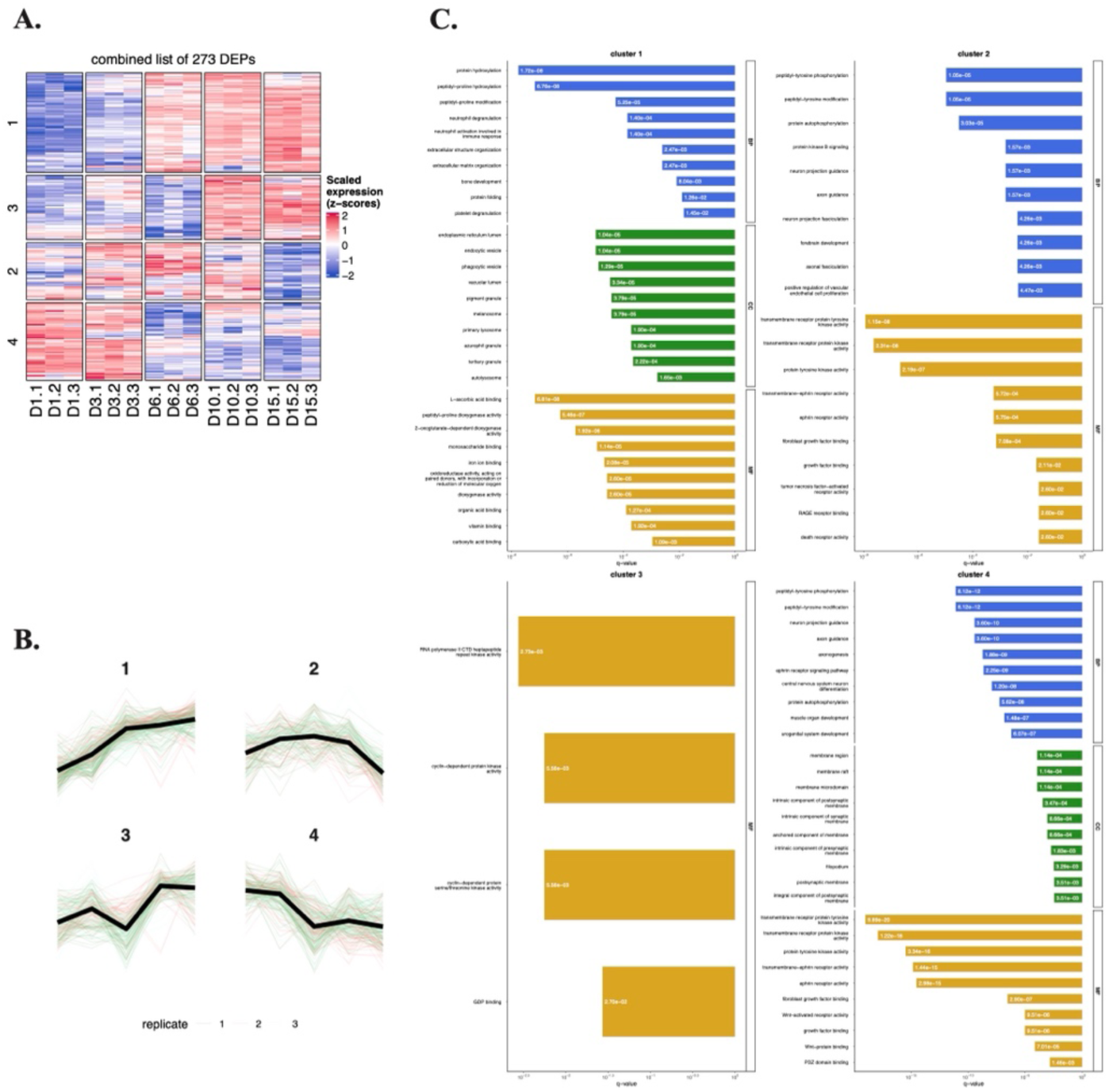
Unsupervised clustering analysis of differentially expressed proteins in the AOB-enriched subproteome during chondrogenesis using the k-means algorithm. A–B. The expression dynamics of each cluster are visible in clusters 1–4. The *x*-axis represents the time points (days of culturing). **C.** Pathway analysis using Gene Ontology (GO) annotation for the 4 clusters. Selected terms for GO biological processes (BP), cellular component (CC), and molecular functions (MF). Barplot shows significant terms, the *x*-axis is gene (protein) ratio and the number of proteins belonging to given enriched terms. Together, these clusters outline the progressive remodeling of the chondrogenic surfaceome.

**Cluster 1** (96 proteins) exhibited progressive upregulation during *in vitro* chondrogenesis, enriched in GO terms such as ECM organization, L-ascorbic acid binding, and peptidyl-prolyl hydroxylation (related to collagen alpha-chain synthesis), and bone development (Figure 5C). In addition to an overrepresentation of collagens, surfaceome cluster 1 contains key chondrocyte regulators and cartilage matrix constituents including MATN1 (pericellular mechanotransduction), SDC4 (syndecan-4; involved in ECM signaling), EPYC (epiphycan; a proteoglycan modulating collagen fibrillogenesis), TRPV4 (transient receptor potential vanilloid 4; involved in mechanosensation), and SFRP3 (secreted frizzled-related protein 3; a Wnt pathway inhibitor), together highlighting its biological significance for cartilage phenotype maintenance and differentiation (42-44).

**Cluster 2** (60 proteins) showed peak expression on culture day 6, with overrepresented GO terms such as kinase activity, ephrin receptor activity, growth factor binding, and FGF signaling. Surfaceome cluster 2 harbors several cell-surface regulators of chondrocyte biology, including RECK (a membrane-anchored matrix metalloprotease inhibitor (45)), FGFR2 (fibroblast growth factor receptor 2; regulates chondrocyte proliferation/differentiation signaling (46)), multiple ephrin receptors (EPHA7, EPHB2, EPHB1, EPHB3 with potential roles in cartilage morphogenesis (47)), SRC kinase (chondrocyte phenotype control (48)), and GPC1 (glypical-1; a proteoglycan and cell surface co-receptor involved in growth factor pathway modulation (49)), highlighting surfaceome mechanisms in cartilage development and regeneration.

**Cluster 3** (63 proteins) displayed transient suppression on culture day 6, followed by late-stage upregulation in mature cultures, with overrepresented GO terms related to RNA polymerase II and the cell cycle. This cluster includes surfaceome and secreted proteins with established contributions to cartilage biology, such as FGFR3 (regulation of proliferation and differentiation (50)), TGFR2 (matrix synthesis and hypertrophy control), TIMP3 (matrix protection), TSP2, and ANXA1 (annexin A1; chondroprotection and matrix homeostasis (51)), together highlighting the complexity of chondrocyte surface signaling and matrix regulation

**Cluster 4** (88 proteins) declined steadily after day 3 in mature chondrocytes, suggesting roles restricted to early differentiation. Enriched GO terms in this cluster include muscle and nervous system development, as well as tyrosine kinase and growth factor signaling. This group included key regulatory surfaceome molecules such as SMO (smoothened; involved in the Indian Hedgehog (IHH) pathway and hypertrophy (52)), or FZD7 (frizzled 7; a Wnt receptor involved in chondroprotection (53)), highlighting the involvement of diverse signaling and adhesion pathways in cartilage biology and prospective OA therapy targets. This cluster contained several proteins without established roles in chondrogenesis, including PODXL (matrix adhesion and phenotype stability) and CNTFR (growth factor signaling and repair).

We also performed a detailed PPI network analysis of each surfaceome cluster using STRING (Figure S4). In **Cluster 1**, the PPI network is enriched in ECM remodeling, collagen biosynthesis, and protein folding, with additional roles in vesicle trafficking and oxidative stress response. The PPI network in **Cluster 2** was dominated by ephrin signaling with four Eph receptors, highlighting roles in developmental patterning. Endocytosis and RAB GTPases are central to membrane dynamics and intracellular transport. SRC and FGFR2 link focal adhesion and growth factor signaling. **Cluster 3** is enriched in cytoskeletal organization, calcium signaling, and cell adhesion pathways. Focal adhesion and actin cytoskeleton pathways dominate the network, driven by actinin, myosin, and dystrophin. Calcium signaling may integrate metabolic and inflammatory responses via CALM3 and S100A6. HMGB1 links nuclear DNA repair with extracellular VEGF signaling. **Cluster 4** is dominated by ephrin signaling with several Eph receptors, highlighting roles in cell migration during nodule formation. Wnt and Hedgehog signaling pathways are central to developmental processes. EGFR/FGFRs link growth factor signaling to MAPK activation.

### 5. Surfaceome enrichment enhances detection of low-abundance targets

Comparative analysis revealed limited overlap (23%) between proteins identified in total lysates and surfaceome-enriched fractions (Table S2). Notably, 77% of AOB-enriched proteins were undetectable in total lysates, demonstrating the method’s enhanced capacity to resolve low-abundance membrane-associated targets. Conventional whole-cell proteomics is constrained by the dominance of high-abundance cytoplasmic and organellar proteins, which often mask low-abundance surface-residing targets during MS analysis. By selectively enriching surface proteins, our approach mitigates this dynamic range limitation, enabling robust identification of membrane-associated and extracellular proteins that otherwise fall below detection thresholds in unfractionated lysates. This methodological advance underscores the efficacy of this method in resolving biologically critical surfaceome components, which are frequently underrepresented in global proteomic analyses, and highlights its utility for advancing receptor, biomarker, and therapeutic target discovery.

### 6. In silico and in vitro chondrogenic surfaceome

To maximize the identification of cell surface proteins, we inferred cell surface localization for all detected proteins using established GO annotations for cellular component and molecular function (16). We first applied this *in silico* prediction on the total cell lysate dataset, defining the GO term-based surfaceome of chondrogenic cultures. Of the 5,207 proteins in this dataset, 885 proteins (17%) were predicted to localize to the cell surface.

We then applied the same approach on the AOB-enriched dataset. The AOB-based enrichment method substantially increased the proportion of identified cell surface proteins, with 55–60% of proteins at all time points and across biological replicates annotated as cell surface-localized (Figure 6A). This represents a significant improvement in the number of surface proteins detected as chondrogenic differentiation progressed, as demonstrated by full symmetrical matrix analysis (Jaccard index) (Figure 6B). We classified the identified surfaceome proteins into major functional categories based on GO molecular function and biological process annotations: enzymes, transporters, receptors, cell junctions and adhesion proteins, ECM components, and unclassified/miscellaneous proteins (including known interacting or binding partners of surface proteins; Table S3). The relative distribution of surface proteins in these categories remained consistent throughout chondrogenesis.

**Figure 6.**
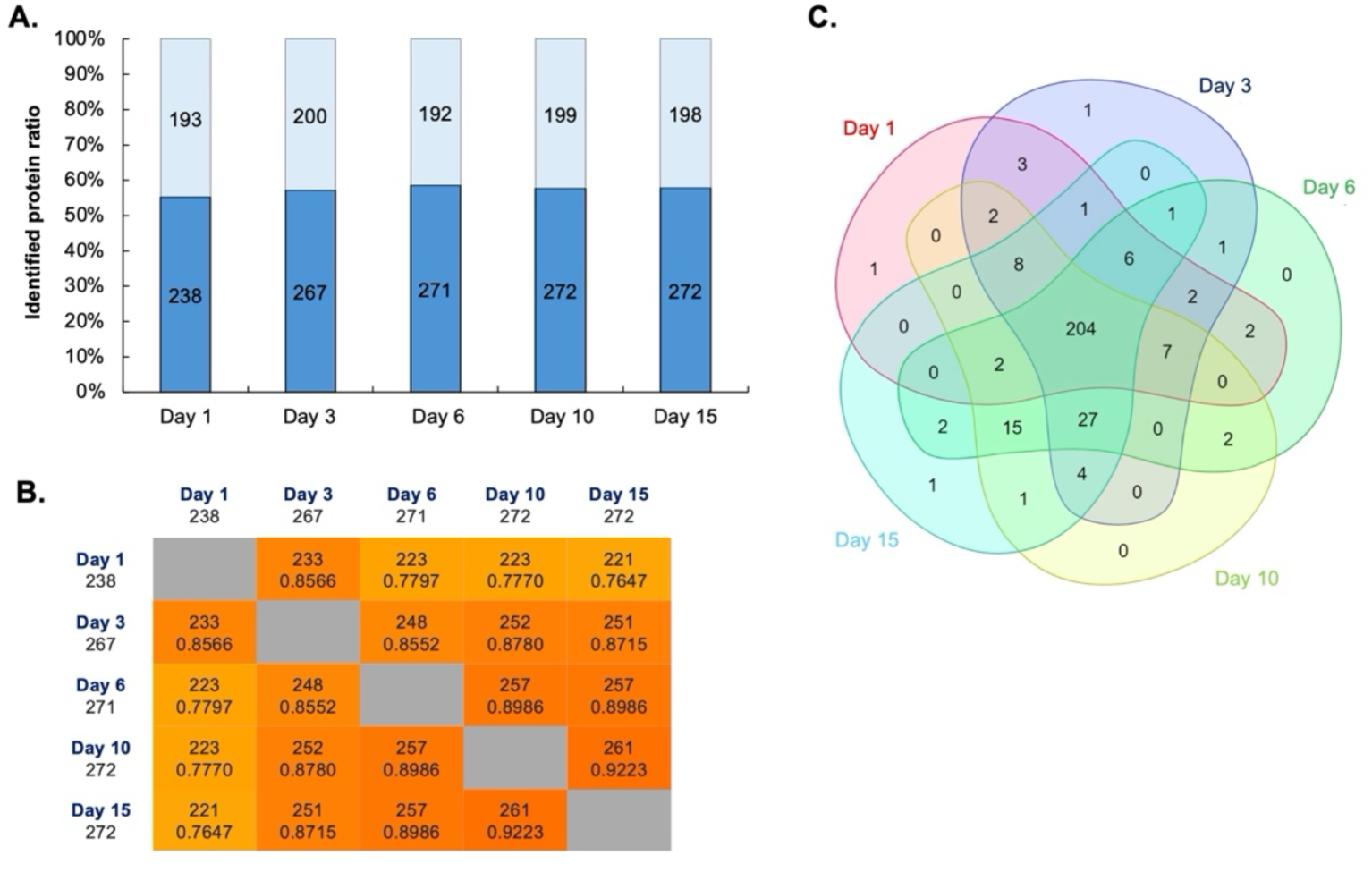
Enrichment of cell surface proteins using the AOB labelling approach. **A.** Distribution of the 504 surface proteins identified in this study (after manual curation using GO terms; dark bars, core surfaceome; light bars, probably non-surface proteins) between different culture days. **B.** Full symmetrical matrix of the surfaceome proteins across differentiating cultures (integers represent the number of surfaceome proteins; non-integers represent Jaccard indices). **C.** Distribution of the manually curated cell surface proteins identified in this study between micromass cultures of various ages.

We next examined qualitative expression dynamics of the cell surface proteins to identify candidate surface biomarkers specific to the early stages of chondrogenic differentiation (days 1–6). Using Venn diagram analysis of manually curated protein lists across the studied time points (Figure 6C), only two proteins met the criteria for early-stage specificity: ciliary neurotrophic factor receptor (CNTFR) and podocalyxin (PODXL).

### 7. Validation of CNTFR and PODXL expression

To verify the expression of CNTFR and PODXL during chondrogenesis, we performed western blot analyses on total cell lysates obtained from micromass cultures harvested at the same time points as AOB-enriched samples. Both CNTFR and PODXL were clearly detected on the blots. For CNTFR, a distinct band was observed at approximately 43 kDa, consistent with its predicted molecular mass, and its expression displayed a gradual decrease over the course of differentiation (Figure 7A). Similarly, PODXL was detected as a prominent ∼60 kDa band, also showing a progressive downregulation throughout the differentiation period (Figure 7B). Notably, the expression patterns of both CNTFR and PODXL closely mirrored the trends observed in our quantitative proteomics analyses performed on the same cultures.

**Figure 7.**
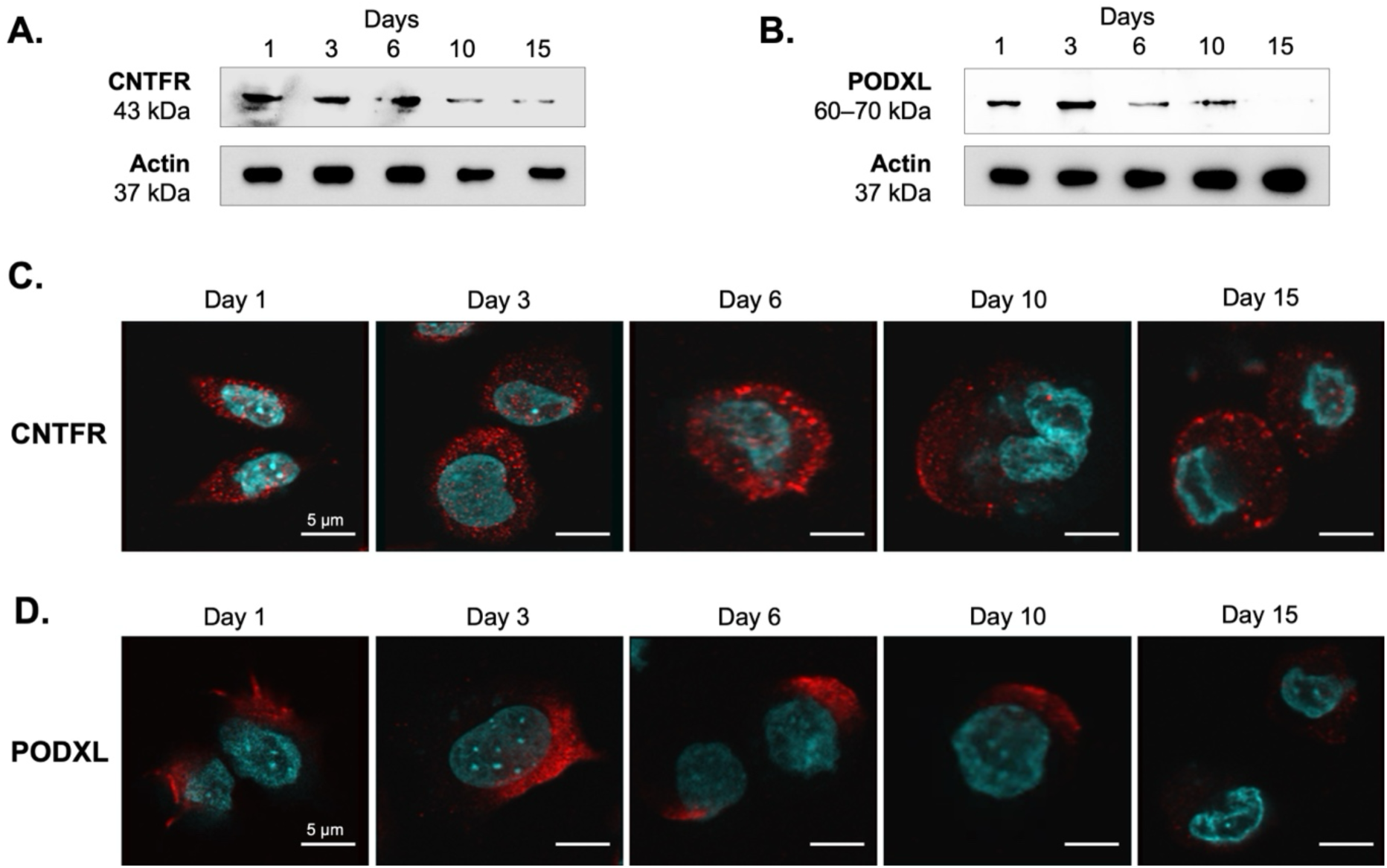
Validation of the efficacy and specificity of the AOB-based glycocapture method. Western blot for CNTFR (**A**) and PODXL (**B**) performed on total cell lysates of micromass cultures. Representative blot image (n = 3). Full-length uncropped blots are presented in Figure S5. Intracellular distribution of CNTFR (**C**) and PODXL (**D**) in cells of micromass cultures detected by immunocytochemistry. Primary antibodies were visualised using Alexa555-conjugated anti-rabbit secondary antibodies (red). Nuclear DNA was stained with DAPI (cyan). Data shown are representative out of 3 independent experiments. Scale bar, 5 µm.

To further investigate the subcellular localization of CNTFR and PODXL, we carried out immunocytochemistry followed by confocal microscopy. Both CNTFR and PODXL were localized to the plasma membrane in chondrogenic cells, with additional intracellular staining (Figure 7C–D). Although this analysis was qualitative, both markers appeared to be downregulated at later stages of differentiation, with this trend being particularly pronounced for PODXL. Notably, PODXL was primarily localized at the leading edge of cellular processes, suggesting a potential role in cell morphology or migration during chondrogenesis.

To investigate the functional role of CNTFR and PODXL in our chondrogenic model, we analyzed their regulatory networks using transcriptomic and proteomic data. STRING-derived interaction network for both proteins (top 25 interactants) were cross-referenced with expression profiles from our previously published RNA-seq dataset (17) of the same chondrogenic model, and total lysate proteomics.

For CNTFR, 21 of the 25 interactants were expressed at the transcript level (Figures S6–S7), and 8 also detected as proteins in the total lysates. PODXL showed expression of 14 interactant transcripts and 8 corresponding proteins. Weighted gene co-expression network analysis (WGCNA) of RNA-seq data, using CNTFR or PODXL expression levels across different samples as a trait, revealed clusters of genes highly correlated with CNTFR or PODXL expression. Therefore, we screened the proximal associated proteins of CNTFR or PODXL and their top 25 interactants within significantly correlated modules and integrated them. The rationale behind this approach was to check the enrichment of GO information in the co-expression modules. For CNTFR, five module eigengenes (MEs) were showing significant degree of correlation (with the turquoise module showing the strongest correlation to the trait), indicating that genes in these MEs may be associated with CNTFR or its top 25 interactants. We therefore performed functional enrichment analysis on these modules, in which GO terms related to tyrosine phosphorylation, cell adhesion, cytokine and growth factor activity, PI3K/Akt and MAPK pathways were enriched. For PODXL, out of five MEs showing high degree of correlation with the gene and its interactants, the black ME displayed the strongest correlation to the trait, in which GO terms such as cell-cell junction, potassium channel activity, differentiation and morphogenesis were over-represented.

### 8. Transient gene silencing of CNTFR and PODXL

To evaluate the functional impact of CNTFR and PODXL on matrix gene regulation, we performed transient siRNA-mediated silencing in primary chondrogenic progenitor cells prior to seeding as micromass cultures (culture day 0). Transfection efficiency and gene silencing were confirmed in 2-day-old cultures (48 hours post-transfection), with *CNTFR* and *PODXL* transcript levels reduced by approximately 57% and 49%, respectively, relative to the non-targeting (NT) control (Figure 8A).

**Figure 8.**
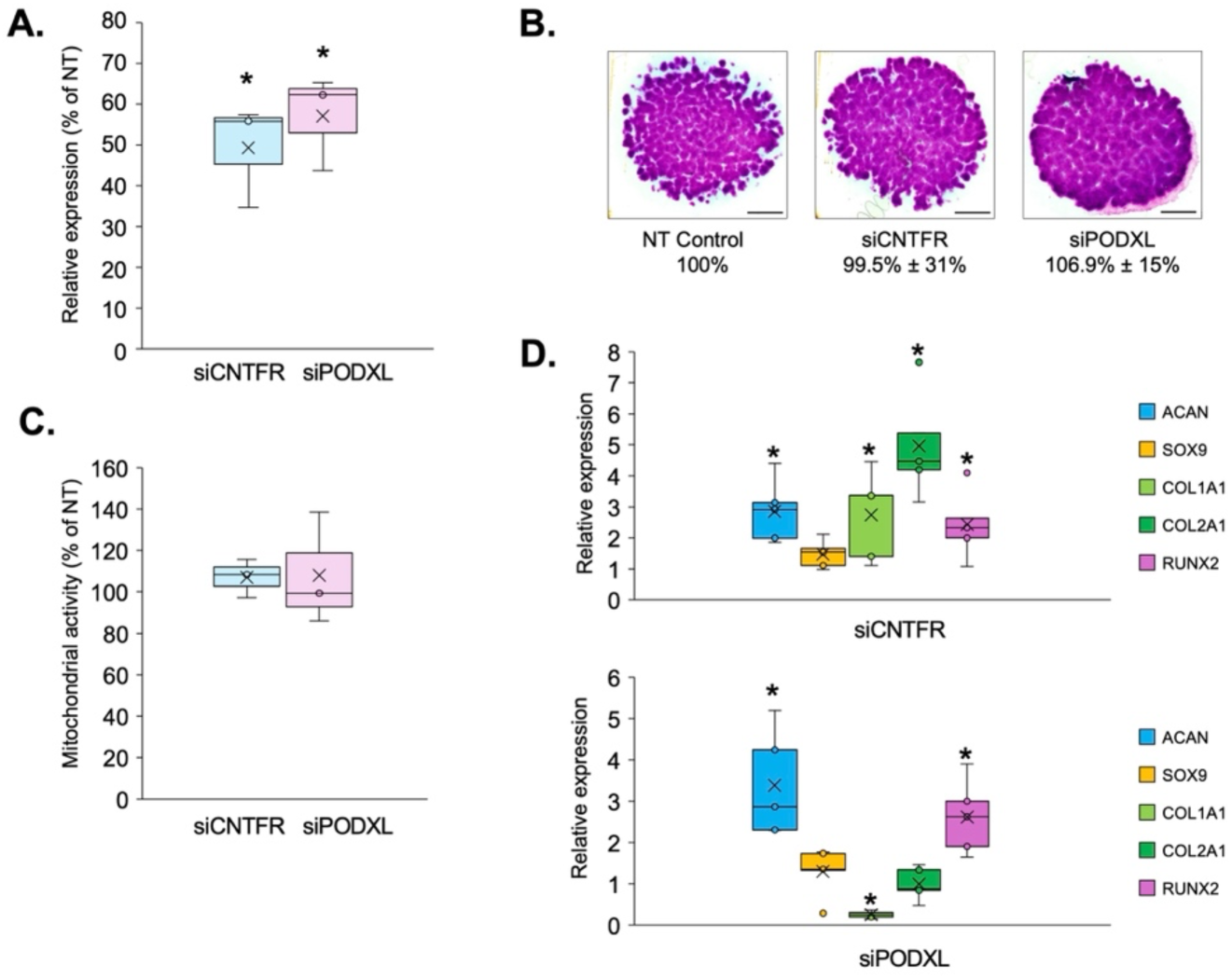
Effects of transient gene silencing of CNTFR or PODXL on osteochondral marker gene expression profiles and cartilage matrix production. **A.** Quantitative RT-qPCR analysis of CNTFR and PODXL mRNA levels in micromass cultures 48 hours post-transfection with target-specific siRNAs. Expression values are shown as fold change relative to non-targeting (NT) control, normalized to the reference gene *RPS7*. Data represent mean ± SEM (n = 3). Asterisks indicate significant differences relative to NT controls (p < 0.05). **B.** Representative DMMB-stained micromass cultures at day 6 illustrating cartilage matrix deposition after transient CNTFR or PODXL silencing compared with NT control. Scale bar = 1 mm. The lower panel shows the corresponding MATLAB-based quantitative analysis of metachromatic area normalized to NT = 100%. Data are mean ± SEM (n = 3). **C.** MTT assay results reflecting relative cell viability in day 6 cultures after silencing. Values are expressed as percentage of NT control (mean ± SEM, n = 3). **D.** Expression of key osteochondrogenic marker genes (*COL1A1, COL2A1, ACAN, RUNX2*) examined 48 hours after transfection. Values represent fold change versus NT control, normalized to *RPS7*. Data are mean ± SEM (n = 3); asterisks indicate p < 0.05.

Although the DMMB-stained micromass cultures (Figure 8B) did not exhibit overt qualitative differences in metachromatic intensity across experimental groups, we additionally performed quantitative image analysis to ensure objective comparison of matrix yield. Using MATLAB-based area quantification normalized to the non-targeting (NT) control (set as 100%), the mean ± SEM values confirmed that total metachromatic matrix deposition remained statistically unchanged following transient knockdown of *CNTFR* or *PODXL* (n = 3). This finding supports the notion that the limited 48-hour window after siRNA treatment is sufficient to capture early transcriptional but not morphological effects, as ECM accumulation in this model typically becomes detectable only several days later. Similarly, cell viability and proliferation remained unaffected on day 6 as assessed by MTT assay (Figure 8C).

However, RT-qPCR analysis performed 48 hours post-transfection revealed distinct transcriptional effects on osteochondrogenic marker gene expression, consistent with early transcriptional rather than morphological responses (Figure 8D). *CNTFR* silencing resulted in marked upregulation of *ACAN* (2.9-fold), *COL2A1* (4.9-fold), *RUNX2* (2.5-fold), and *COL1A1* (2.7-fold) relative to non-targeting (NT) control cultures, whereas *PODXL* silencing increased *ACAN* (3.3-fold) and *RUNX2* (2.6-fold) expression but downregulated *COL1A1* (–3.8-fold). These divergent transcriptional responses indicate that *CNTFR* and *PODXL* differentially modulate ECM-related gene networks, potentially influencing the matrix composition rather than its total production.

Although neither perturbation affected observable changes in ECM deposition or proliferation, these findings highlight distinct effects on transcriptional regulation of ECM-associated genes by *CNTFR* and *PODXL*. The contrasting modulation of *COL1A1* suggests that these surface proteins may influence the qualitative composition and molecular specificity of the ECM during chondrogenesis. Furthermore, the observed expression patterns reinforce the concept that *PODXL* and *CNTFR* act as early regulatory nodes, modulating transcriptional programs that precede measurable ECM remodeling during chondrogenesis.

## Discussion

In this work, we identify podocalyxin (PODXL) and ciliary neurotrophic factor receptor (CNTFR) as previously unrecognized cell surface constituents of chondrogenic progenitor cells. By employing a glycocapture-assisted subproteome enrichment strategy combined with high-throughput shotgun mass spectrometry, we achieved enhanced sensitivity for detecting low-abundance surface compared to conventional whole-cell and surfaceome profiling proteomics approaches, as demonstrated in earlier studies (10, 16, 54). While similar enrichment and mass spectrometry-based workflows have previously been applied to characterize the surfaceome of mesenchymal cells and primary equine chondrocytes under normal and pro-inflammatory conditions (9, 15), the present work complements and extends this methodology to systematically and quantitatively profile dynamic changes in the surfaceome during *in vitro* chondrogenic differentiation in embryonic progenitor cells. Our study offers a temporal and functional proteomic map of the chondrogenic cell surface in a developmental (non-inflammatory) context, widening the molecular framework for understanding how membrane proteins contribute to cartilage lineage specification and phenotype stability.

### CNTFR and implications for chondrogenesis

The identification of CNTFR in differentiating chondrocytes opens new perspectives on neurotrophic cytokine signaling in cartilage biology. Ciliary neurotrophic factor (CNTF), a member of the interleukin-6 (IL-6) family, signals through a tripartite receptor complex comprising CNTFRα, gp130, and LIFR (55, 56). Initially characterized as a neuronal survival and differentiation factor, CNTF has since been implicated in multiple tissues, including bone and cartilage (57-59). Related ligands such as cardiotrophin-like cytokine (CLCF1) also influence MSC differentiation and osteogenesis (60), reinforcing the idea of a functional role for CNTF/CNTFR signaling in skeletal development (61).

CNTFR expression has known roles in cartilage homeostasis and OA progression. For instance, upregulation of the CRLF1/CLCF1 complex, which competes with CNTF for CNTFR binding, disrupts cartilage ECM homeostasis and exacerbates OA (58). Our finding of persistent CNTFR expression during chondrogenesis, coupled with its plasma membrane localization in differentiating chondrocytes, suggests dual roles in cartilage maintenance and pathological remodeling. This positions CNTFR as both a mechanistic regulator and potential therapeutic target in OA.

Our observation aligns with previous findings on signaling-driven regulation of chondrogenic differentiation. In particular, comprehensive analyses of cartilage lineage specification (62) and recent single-cell and transcriptomic profiling of chondrocyte subpopulations (8) highlighted the interplay between growth factor–mediated pathways (including IL-6 family cytokines, FGFR, and TGF-β signaling) and ECM remodeling as key determinants of progenitor maintenance and hypertrophic transition. Our results suggest that CNTFR activity may function within this network, modulating survival and fibrocartilaginous remodeling responses during differentiation.

### Podocalyxin and progenitor state regulation

Podocalyxin (PODXL), classically associated with kidney podocytes and vascular endothelium, is a member of the CD34 transmembrane sialomucin family (63). It contributes to cell adhesion, migration, and progenitor cell maintenance (64, 65). Its presence in chondrogenic cells is novel and suggests potential roles in modulating progenitor cell responses to chondrogenic cues such as SOX9 and TGF-β signaling (66). This is conceptually consistent with previous studies demonstrating dynamic modulation of cell-matrix receptors and cytoskeletal regulators during the dedifferentiation-redifferentiation cycle of chondrocytes (62). Moreover, at the single-cell level, progenitor-like clusters characterized by strong expression of adhesion and polarity molecules were described (8), a feature reminiscent of PODXL-positive states in our study. Together, these findings support the hypothesis that PODXL may contribute to the maintenance of a transitional chondrogenic phenotype with migratory or matrix-adaptation potential. While evidence directly linking PODXL to chondrogenesis is limited, its downregulation during osteogenic maturation and association with progenitor-rich MSC populations highlight it as a candidate regulator of early cartilage differentiation.

### Opposing roles in ECM quality control

Our transient knockdown assays revealed opposing transcriptional regulation of *COL1A1*, a well-known marker of fibrocartilage and mechanically compromised repair tissue (62, 67), by *PODXL* and *CNTFR*. *CNTFR* silencing increased *COL1A1* expression (2.7-fold), whereas *PODXL* silencing reduced it (−3.8-fold). The differential regulation of *COL1A1* between the two knockdowns suggests that these proteins influence ECM-related transcription through distinct pathways. PODXL may help sustain fibrogenic signaling (via β1-integrin/FAK pathways) (68, 69), while CNTFR loss may mimic cytokine deprivation stress, and trigger compensatory fibrotic responses, possibly through ERK/STAT3 signaling cross-talk, reminiscent of synovial fibrosis in OA via impaired signaling (70). Thus, the responses observed here are consistent with the predicted biological roles of CNTFR and PODXL, which are reflected in the network associations shown in Figure S7.

The absence of overt changes in ECM production in our knockdown model suggests functional redundancy in matrix remodeling, post-transcriptional regulation, or compensation by other surface receptors. Quantitative evaluation of DMMB-positive areas using MATLAB-based image analysis further confirmed that total matrix deposition was not significantly altered by either treatment. The absence of visible or quantitative differences therefore likely reflects the temporal separation between transcriptional and phenotypic responses, rather than true lack of functional relevance.

### Translational implications

Single-cell RNA-seq analysis of human articular chondrocytes revealed detectable transcripts for both *CNTFR* (predominantly in effector and homeostatic chondrocytes) and *PODXL* (at lower levels in *VEGFA*^+^ cells, fibrochondrocytes, and pre-hypertrophic chondrocytes) (Figure S8). This, combined with their extracellular accessibility, positions CNTFR and PODXL as potentially detectable biomarkers in synovial fluid or serum, opening avenues for early OA diagnosis and therapeutic monitoring.

Beyond their diagnostic potential, modulating PODXL-mediated cell-matrix interactions or CNTFR-dependent signaling could open new, targeted approaches to cartilage repair or OA management (71). While functional redundancy in chondrogenic signaling networks or compensatory mechanisms from other surface receptors may mask direct phenotypic impact in single knockdowns, combinatorial gene perturbations or pathway-focused interventions could reveal their full regulatory capacity.

### Limitations

The use of chicken embryonic limb bud-derived chondrogenic progenitor cells provides a valuable developmental model but may not fully recapitulate all aspects of human chondrogenesis or OA pathophysiology. While the embryonic avian limb model has been extensively validated as a mechanistic system for studying cartilage differentiation, species-specific variations in ECM composition, signaling dynamics, and progenitor populations may limit the direct extrapolation of our findings to human articular cartilage. Similarly, the micromass culture system, although well-established for modeling chondrogenic lineage commitment (18), remains a simplified *in vitro* system that cannot reproduce the full cellular complexity, mechanical stimuli, or biochemical heterogeneity present in the native joint environment.

Our cell surface protein enrichment workflow focused on N-linked glycosylated plasma membrane constituents, which ensures strong selectivity for *bona fide* surface proteins but may underrepresent other biologically important groups such as non-glycosylated or O-linked membrane proteins and secreted matrix components. Consequently, certain signaling or adhesion receptor families involved in chondrogenic regulation may not have been captured. In addition, although Gene Ontology (GO) annotation was performed using human ortholog mapping to facilitate cross-species interpretation, specific biological processes may be unique to avian systems or not entirely conserved between chicken and human cartilage (72, 73).

In addition, functional redundancy among surface receptors may partially mask morphological or matrix-level phenotypes, as parallel signaling pathways can compensate for transient loss of individual components. The transient siRNA-based functional assays for *PODXL* and *CNTFR* were intentionally designed to capture short-term transcriptional responses; however, such brief modulation may not fully reveal cumulative effects on long-term ECM assembly, organization, or mechanical integrity.

Future studies incorporating human chondrogenic progenitors or patient-derived chondrocyte models, coupled with more comprehensive surfaceome coverage and stable gene perturbation strategies, will be necessary to confirm the translational relevance and fully delineate the downstream signaling networks of these proteins and confirm the translational relevance of the observed regulatory patterns. Fully comprehensive surfaceome profiling of chondrogenic and mature chondrocytes will ultimately require integration of alternative enrichment chemistries, spatial proteomics, and cell types, as well as deeper sampling under both physiological and disease-mimicking conditions.

## Conclusions

In summary, we map the dynamic surfaceome of chondrogenic cells and identify PODXL and CNTFR as newly identified components at the interface of cell communication, adhesion, and survival signaling, showing expression dynamics consistent with roles in ECM regulation. These findings deepen our understanding of chondrocyte surface signaling networks, hold promise for biomarker development and could inform therapeutic strategies in OA and cartilage regeneration.

Our study contributes new insight into surfaceome remodeling during embryonic chondrogenesis. The candidate surface proteins identified here provide a basis for further discovery efforts and functional exploration under a wider range of conditions. Future studies in human cellular and disease models will be crucial to determine their diagnostic and therapeutic potential. Furthermore, follow-up studies will be needed to expand the surfaceome landscape under diverse physiological (e.g., hormonally regulated, loaded cartilage) and pathological (inflammatory, OA-like) conditions. The dynamic and context-dependent nature of the chondrocyte surfaceome warrants integration of multiple complementary strategies, including alternative cell models, stimuli, and orthogonal enrichment techniques to approach a more complete characterization.

## Additional files (Supplementary material)

- Additional file 1 (PDF) contains Supplementary Figures S1–S8 and Supplementary Tables S1–S3, providing additional data supporting the main findings.
- Additional files 2–5 (Excel spreadsheets) include: differential expression analysis and clustering results for total lysate proteomes (files 2–3) and surfaceome proteomes (files 4–5). Collectively, these materials offer expanded experimental details, data visualizations, and full quantitative proteomic datasets underlying the study.

## DECLARATIONS

### Ethics approval and consent to participate

Not applicable to thus manuscript.

### Consent for publication

Not applicable to thus manuscript.

### Availability of data and materials

The dataset supporting the conclusions of this article is available in the ProteomeXchange Consortium *via* the PRIDE partner repository with the dataset identifier PXD064013 (https://www.ebi.ac.uk/pride/archive/projects/PXD064013/private)

### Authors’ contributions

All authors made substantial contributions to discussion of the content, writing of the original outline, and reviewing/editing of the article before submission. All authors approve the final version for publication and agree to be accountable for the accuracy and integrity of the study. Patrik Kovács: Formal analysis, Methodology, Validation, Visualisation, Writing—original draft, Writing—review & editing. Peter Brazda: Formal analysis, Visualisation, Writing—original draft, Writing—review & editing. Tibor Hajdú: Formal analysis, Methodology, Writing—original draft, Writing—review & editing. Boglárka Harsányi: Formal analysis, Methodology, Writing— original draft. Krisztián Juhász: Formal analysis, Methodology, Writing—original draft. Roland Takács: Formal analysis, Methodology, Validation, Writing—original draft, Writing—review & editing. Judit Vágó: Formal analysis, Methodology, Writing—original draft, Writing—review & editing. Zhangzgeng Wang: Formal analysis, Methodology, Writing—original draft, Writing— review & editing. Clare Coveney: Formal analysis, Methodology, Writing—original draft, Writing—review & editing. David J. Boocock: Formal analysis, Software, Methodology, Visualization, Writing—original draft, Writing—review & editing. Csaba Matta: Conceptualization, Formal analysis, Funding acquisition, Methodology, Supervision, Validation, Writing—original draft, Writing—review & editing.

## Acknowledgements

The authors thank Mrs. Krisztina Biróné Barna and Ms. Éva Katona from the Department of Anatomy, Histology and Embryology, and Prof. Éva Csősz and Mrs. Julianna Kökényesiné Csáki from the Proteomics Core Facility, Department of Biochemistry and Molecular Biology, Faculty of Medicine, University of Debrecen, for technical assistance.

