## Supplementary figures S1-S8; Supplementary tables S1-S3. for "Podocalyxin and ciliary neurotrophic factor receptor are novel components of the surfaceome of chondrogenic cells"

### Contents

| | |
|---|---|
| *Figure S1.* | Volcano plots showing differentially expressed proteins (DEPs) between consecutive time points, and between undifferentiated (day 1) and mature/hypertrophic (day 15) chondrogenic micromass cultures (total lysates). |
| *Figure S2.* | Protein-protein interaction (PPI) analysis of the 6 clusters in total cell lysates using STRING based on the top 20 (hub) proteins. In each network, the size of the circles represented the degree of value: larger circles represent proteins with more connections (higher degree), meaning they interact with more other proteins in the network. |
| *Figure S3.* | Volcano plot highlighting differentially expressed proteins between consecutive time points, and between Day 1 (undifferentiated) and Day 15 (mature/hypertrophic) chondrogenic micromass cultures after AOB-enrichment. |
| *Figure S4.* | Protein-protein interaction (PPI) analysis of the 4 clusters in surfaceome samples using STRING. In each network, the size of the circles represented the degree of value: larger circles represent proteins with more connections (higher degree), meaning they interact with more other proteins in the network. |
| *Figure S5.* | Full-length uncropped blots for the western blot images presented in Figure 7. |
| *Figure S6.* | Weighted gene co-expression network analysis (WGCNA) results performed on the NGS dataset to identify clusters of highly correlated genes to CNTFR or PODXL, using culture age (time) as a trait. |
| *Figure S7.* | Transcript and protein expression patterns for the closest 25 interactants, as well as WGCNA co-expression networks, and enriched gene ontology (GO) terms in the MEs for CNTFR (A) and PODXL (B). |
| *Figure S8.* | In a single-cell atlas of healthy cartilage tissue, we identified eight chondrocyte clusters and systematically annotated their cellular identities (upper panel). Subsequent analysis revealed the expression patterns of CNTFR and PODXL across these chondrocyte subpopulations (lower panel). |
| *Table S1.* | Forward and reverse primer sequences, gene accession numbers and amplicon length for each primer pair employed in this study. |
| *Table S2.* | Comparative analysis of all total lysate and surfaceome-enriched samples, which revealed limited overlap, with only approx. 30% of proteins identified in both fractions. |
| *Table S3.* | Relative distribution of GO functional classification of the identified surface proteins during chondrogenic differentiation in micromass cultures. |

*Figure S1.* Volcano plots showing differentially expressed proteins (DEPs) between consecutive time points, and between undifferentiated (day 1) and mature/hypertrophic (day 15) chondrogenic micromass cultures (total lysates). Cutoff values: $\log_2$ fold change > 1 (*x*-axis) and adjusted *p*-value < 0.05 (*y*-axis). Red dots signify proteins that were upregulated in later time points, whereas blue signifies proteins downregulated in later time points.

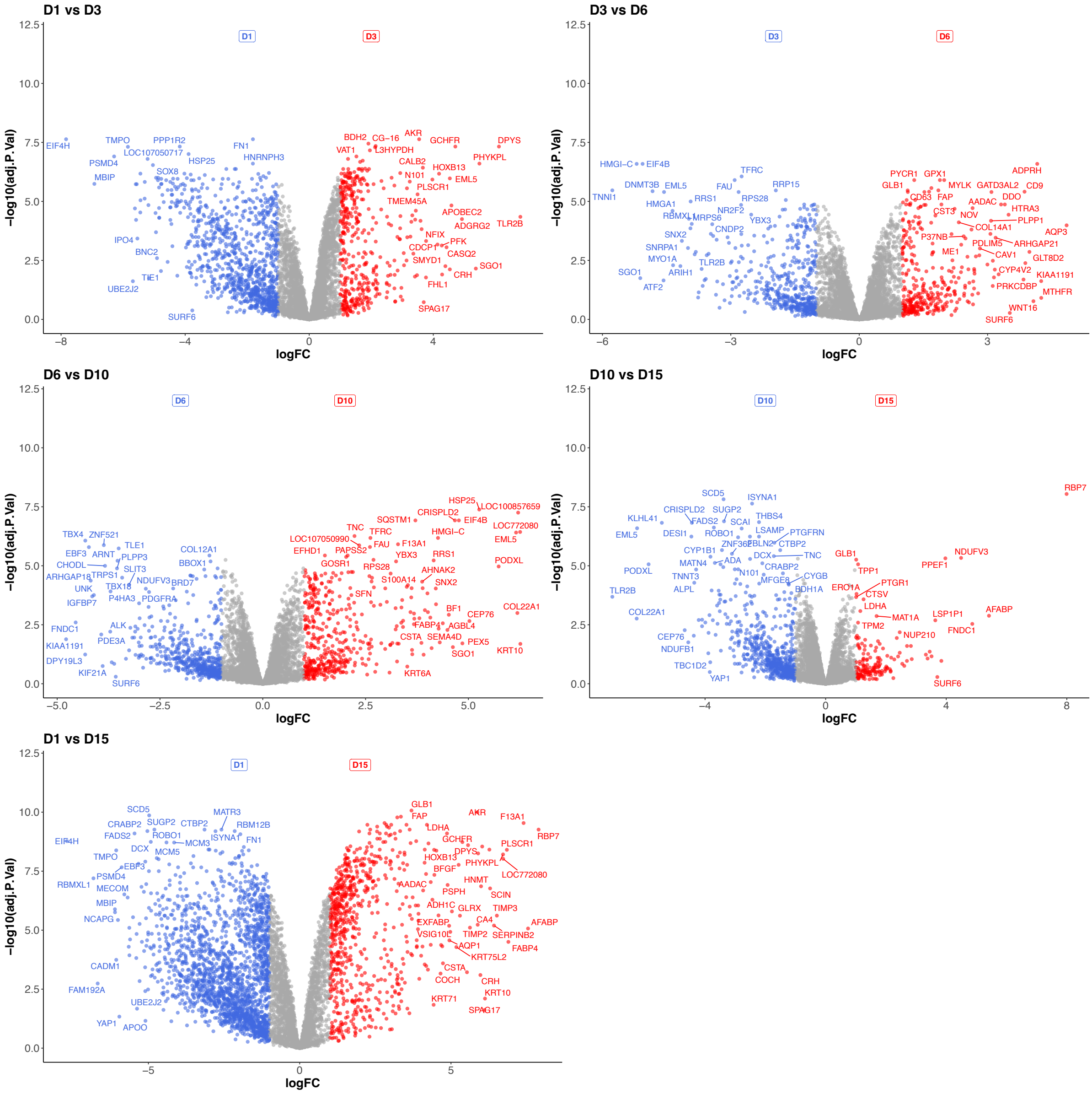

*Figure S2.* Protein-protein interaction (PPI) analysis of the 6 clusters in total cell lysates using STRING based on the top 20 (hub) proteins. In each subnetwork, the size of the circles represented the degree of value: larger circles represent proteins with more connections (higher degree), meaning they interact with more other proteins in the network.

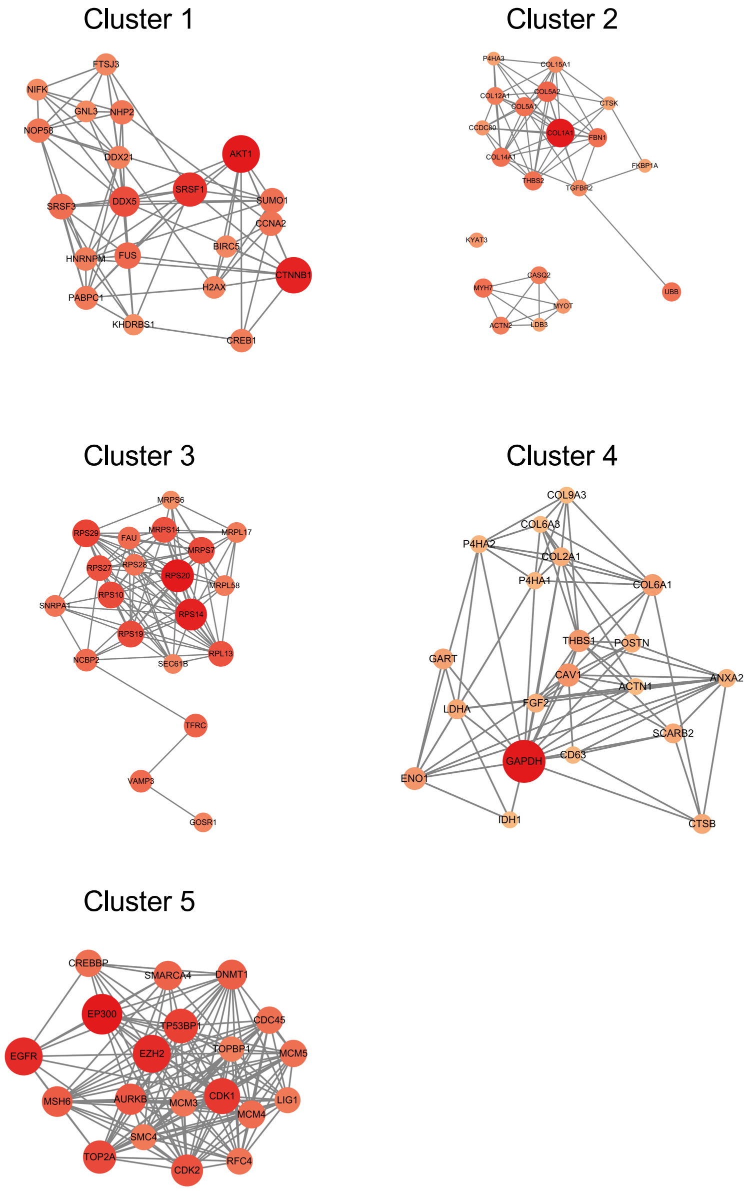

*Figure S3.*Volcano plot highlighting differentially expressed proteins between consecutive time points, and between Day 1 (undifferentiated) and Day 15 (mature/hypertrophic) chondrogenic micromass cultures after AOB-enrichment. Cutoff values: $log_2$ fold change > 1 (horizontal axis) and adjusted *p*-value < 0.05 (vertical axis). Red dots signify proteins that were upregulated in later time points, whereas blue signifies proteins downregulated in later time points.

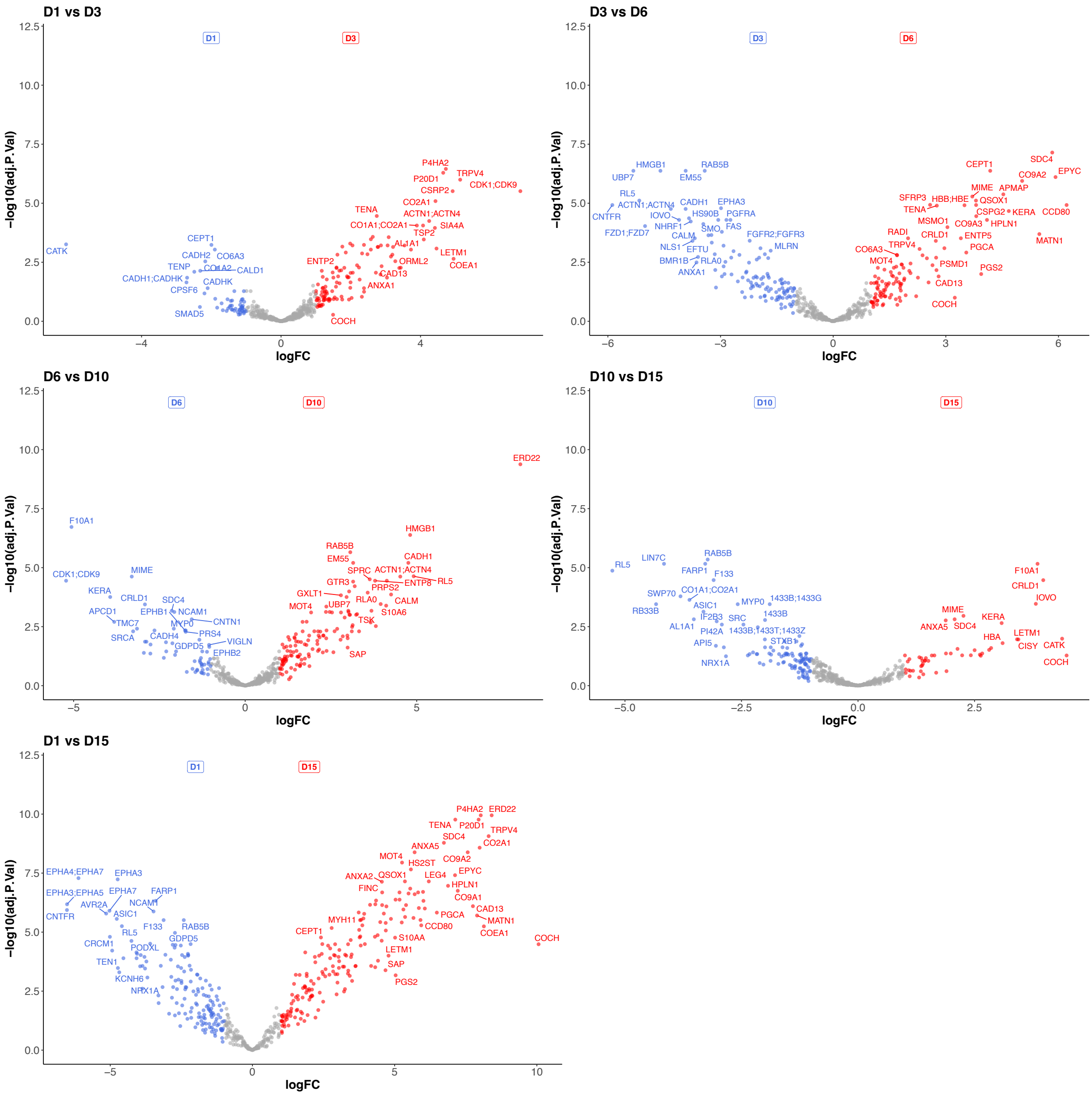

*Figure S4.* Protein-protein interaction (PPI) analysis of the 4 clusters in surfaceome samples using STRING. In each network/subnetwork, the size of the circles represented the degree of value: larger circles represent proteins with more connections (higher degree), meaning they interact with more other proteins in the network.

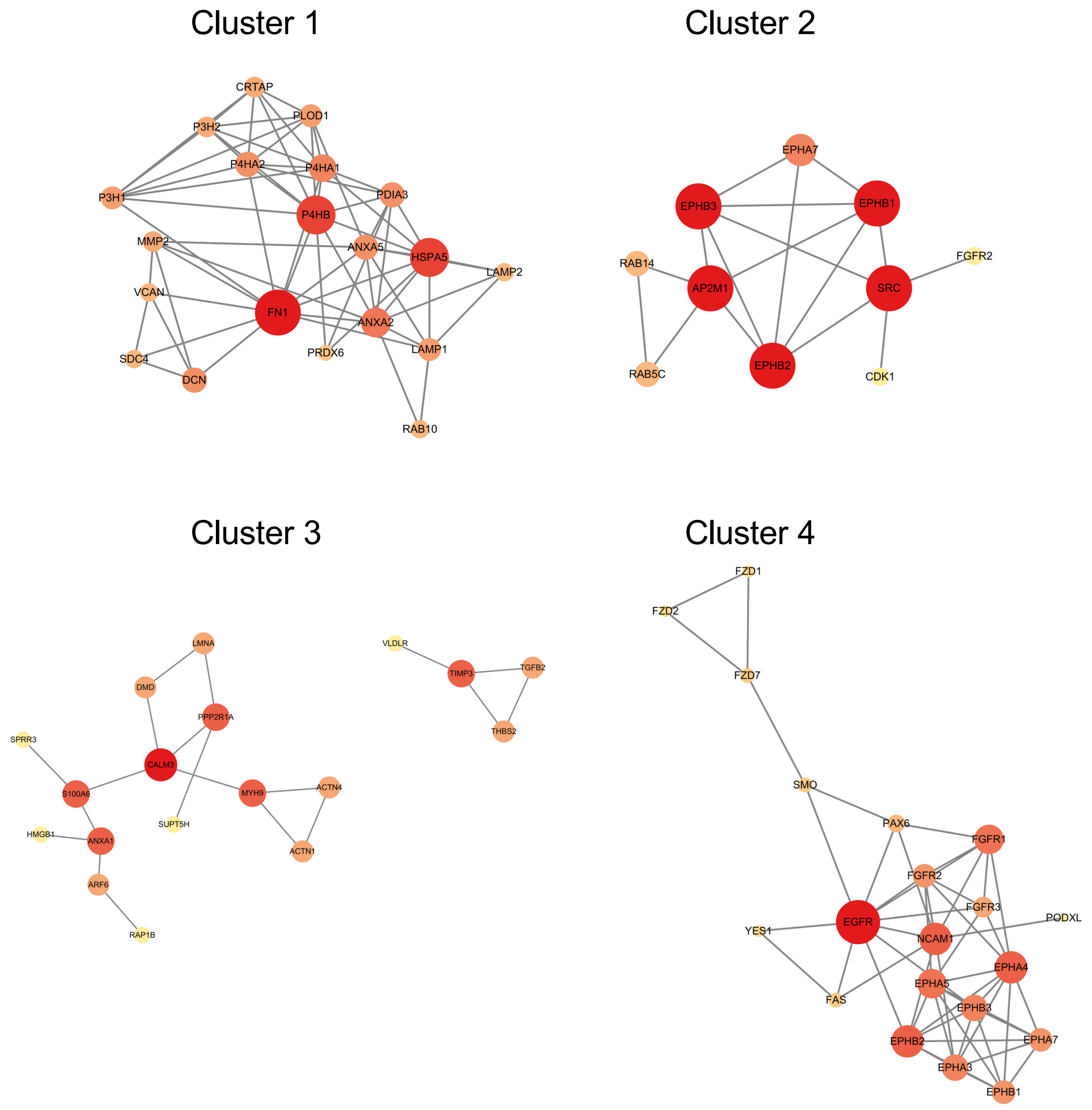

*Figure S5.* Full-length uncropped blots for the western blot images presented in Figure 7.

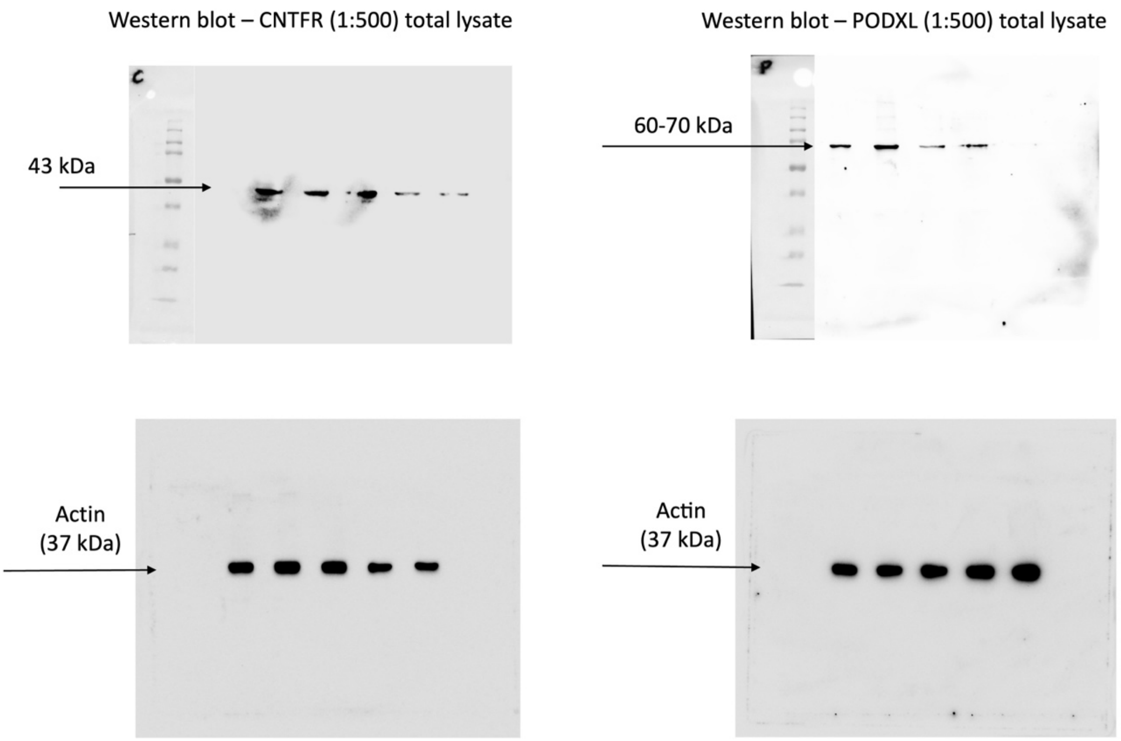

*Figure S6.* Weighted gene co-expression network analysis (WGCNA) results performed on the NGS dataset to identify clusters of highly correlated genes to CNTFR or PODXL, using culture age (time) as a trait. For CNTFR, the turquoise module eigengene (ME) was showing the highest degree of correlation; for PODXL, the black ME showed the highest correlation to the trait.

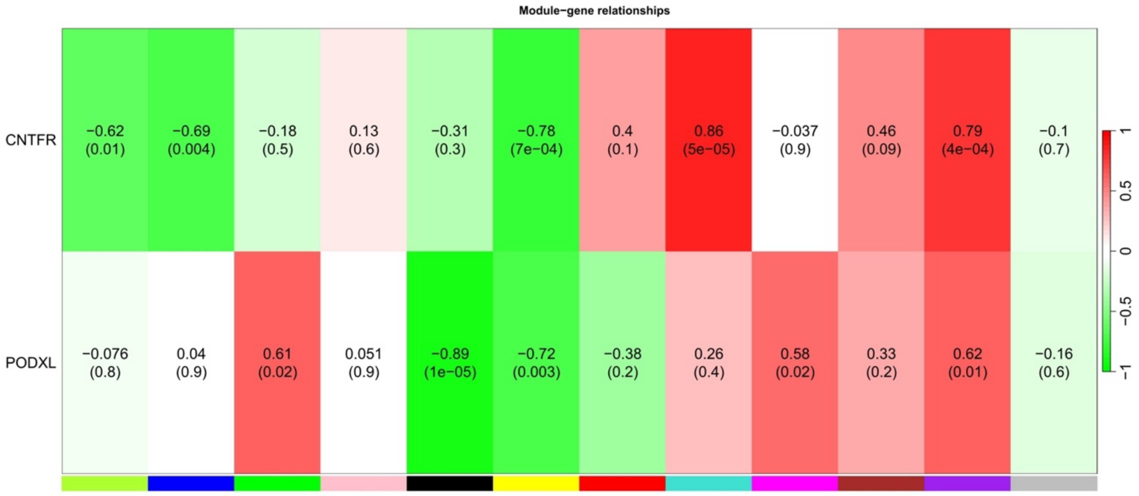


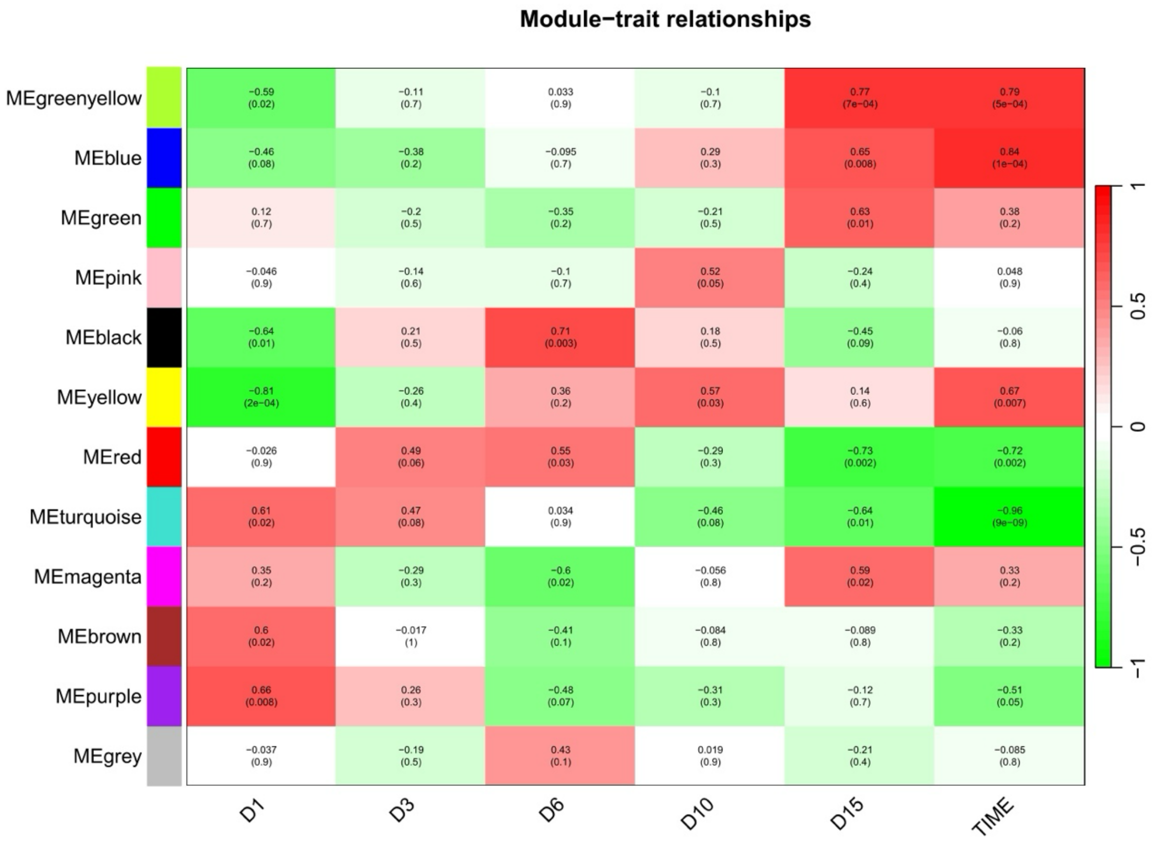

*Figure S7.* Transcript and protein expression patterns for the closest 25 interactants, as well as WGCNA co-expression networks, and enriched gene ontology (GO) terms in the MEs for CNTFR (A) and PODXL (B).

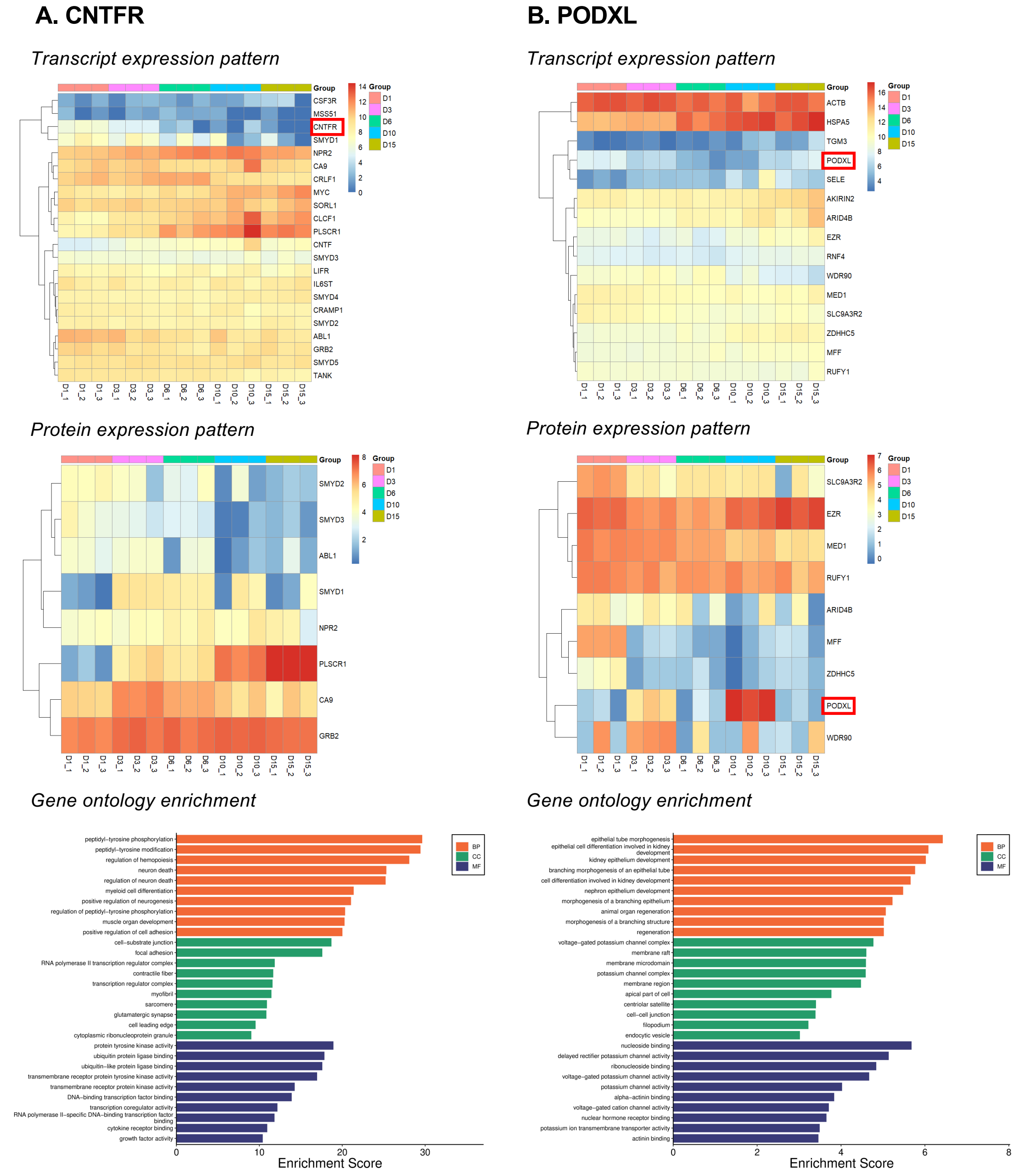

*Figure S8.* The single-cell atlas of healthy cartilage tissue integrates data from two public datasets: cartilage samples derived from the ankle joints of five healthy individuals (GSE216578), and healthy knee joint cartilage specimens obtained from three individuals in dataset GSE255460. Data integration and analytical processing were performed using the Seurat R package. In the single-cell atlas of healthy cartilage tissue, we identified eight chondrocyte clusters and systematically annotated their cellular identities (upper panel). Subsequent analysis revealed the expression patterns of CNTFR and PODXL across these chondrocyte subpopulations (lower panel). Notably, CNTFR demonstrated predominant expression enrichment in both effector chondrocytes (EC) and homeostatic chondrocytes (HomC). This expression signature suggests potential functional regulation of CNTFR during chondrocyte differentiation stages. PODXL demonstrated much weaker expression profile throughout chondrocyte populations, perhaps with somewhat higher transcript levels in pre-hypertrophic chondrocyte (Pre-HTC) subpopulations.

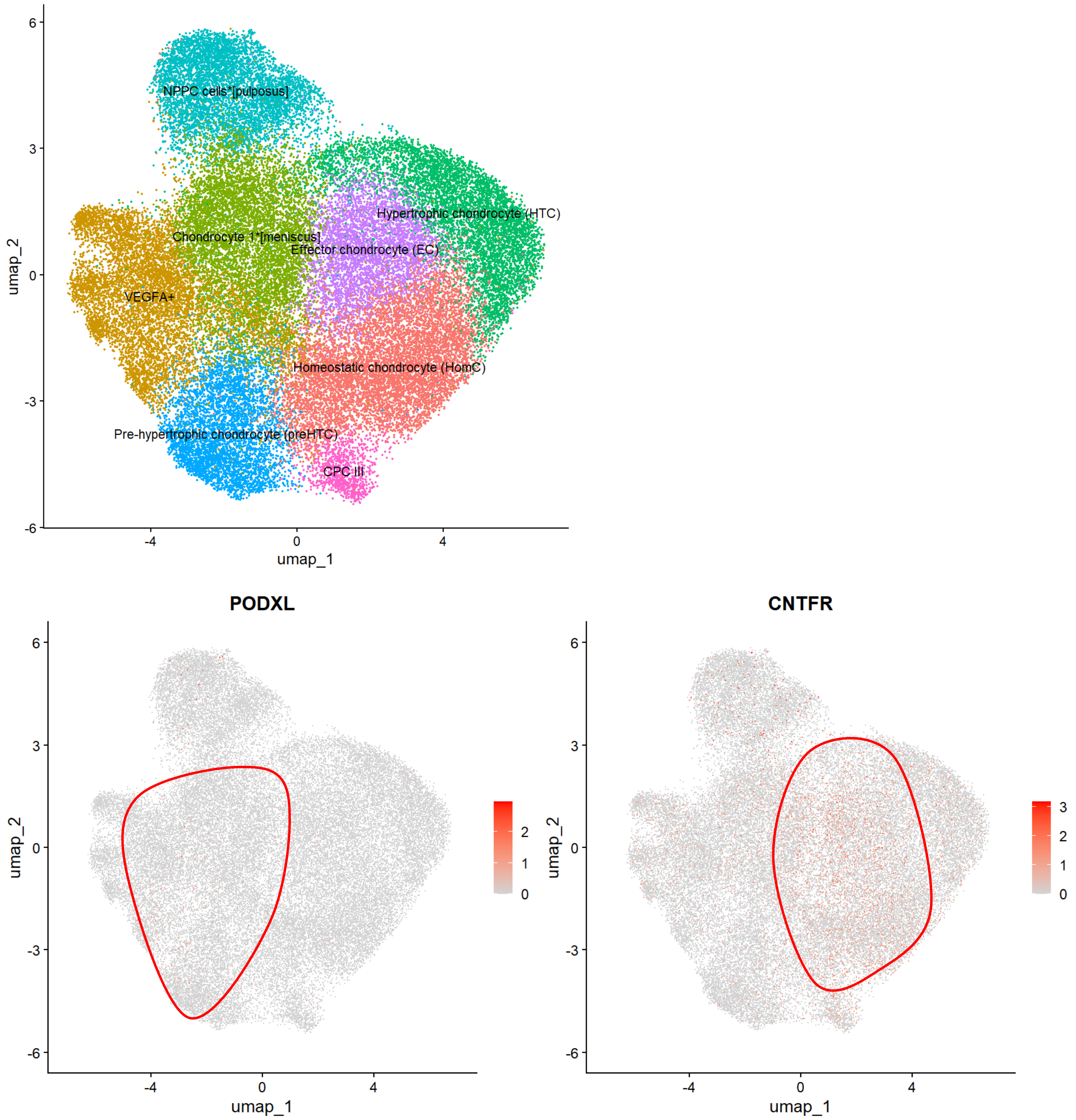


The dot plot below shows that CNTFR is highly expressed mainly in Homeostatic chondrocytes (HomC) and Effector chondrocytes (EC), while PODXL is predominantly expressed in Regulatory chondrocytes (RegC) and Pre-hypertrophic chondrocytes (preHTC).

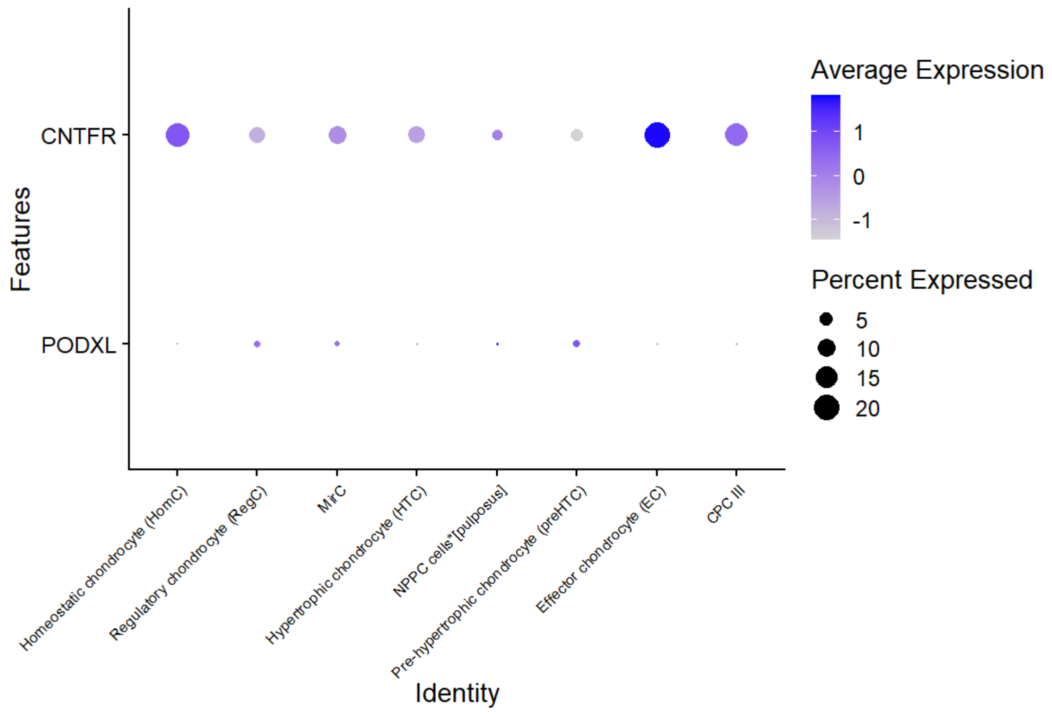
Average Expression
1
0
-1
Percent Expressed
5
10
15
20
CNTFR
PODXL
Features
Homeostatic chondrocyte (HomC)
Regulatory chondrocyte (RegC)
MirC
Hypertrophic chondrocyte (HTC)
NPPC cells*[pulposus]
Pre-hypertrophic chondrocyte (preHTC)
Effector chondrocyte (EC)
CPC III
Identity

*Table S1.* Forward and reverse primer sequences, gene accession numbers and amplicon length for each primer pair employed in this study.

| Gene symbol, species | Accession number | Primer sequence | Product length (bp) |
|---|---|---|---|
| ***RPS7*** *(Gallus gallus)* | XM_004940515.4 | FW: CGGAGTGCCGCGAAAGG<br>REV: TGTGATGTTCAACTCCCGCA | 157 |
| ***YWHAZ*** *(Gallus gallus)* | NM_001031343 | FW: GTTCCCTTGCAAAAACGGCT<br>REV: GAGGCAGACGGAAGTTGGAA | 199 |
| ***PPIA*** *(Gallus gallus)* | NM_001166326.1 | FW: GAGCTCTTCGCTGACAAGGT<br>REV: GCGTAAAGTCACCACCCTGA | 139 |
| ***HPRT1*** *(Gallus gallus)* | NM_204848.1 | FW: TGGTGGGGATGACCTCTCAA<br>REV: TCCAACAAAGTCTGGCCGAT | 190 |
| ***RPL4*** *(Gallus gallus)* | NM_001007479 | FW: TGTTTGCCCCAACCAAGACT<br>REV: CTCCTCAATGCGGTGACCTT | 137 |
| ***RPL13*** *(Gallus gallus)* | NM_204999.1 | FW: GGCCCGTGTTATCTCAGAGG<br>REV: CCGCTTCTTTGGCACGTTTT | 113 |
| ***SOX9*** *(Gallus gallus)* | NM_204281.1 | FW: TTTCCGAGACGTGGACATCG<br>REV: GTACCGCTGTAGGTGGTGAC | 150 |
| ***ACAN*** *(Gallus gallus)* | NM_204955 | FW: AGCAGTAGATGCACTGGGAC<br>REV: GCCAGGTCGATCTCACACAG | 153 |
| ***COL2A1*** *(Gallus gallus)* | NM_204426.1 | FW: GGGACCTCAAGGCAAAGTCG<br>REV: TTCCAGGCTCACCATTAGCG | 140 |
| ***COL1A1*** *(Gallus gallus)* | NM_001396622.1 | FW: TACTGCAACATGGAGACGGG<br>REV: CCGCCGTACTCAAACTGGAA | 152 |
| ***RUNX2*** *(Gallus gallus)* | NM_204128 | FW: GAGTCAGATTACAGACCCCAGG<br>REV: AAATGGGCCCAGCTCGGAA | 199 |
| ***CNTFR*** *(Gallus gallus)* | XM_046934809.1 | FW: ATCACGGATGCCTATGCTGG<br>REV: GTGCTCGTCGTCTCTGTGAT | 164 |
| ***PODXL*** *(Gallus gallus)* | XM_046906716.1 | FW: CATGTCATTGTGCATCGCGT<br>REV: TAGGTGACGTTGGTGATGCC | 101 |

*Table S2.* Comparative analysis of all total lysate and surfaceome-enriched samples, which revealed limited overlap, with only approx. 30% of proteins identified in both fractions.

| Sample ID | Surfaceome only | Total lysate only | Overlap | Overlap % |
|---|---|---|---|---|
| D1_1 | 289 | 5016 | 84 | 22.52% |
| D1_2 | 305 | 5010 | 87 | 22.19% |
| D1_3 | 312 | 5013 | 86 | 21.61% |
| D3_1 | 317 | 4944 | 93 | 22.68% |
| D3_2 | 335 | 4995 | 100 | 22.99% |
| D3_3 | 328 | 4998 | 98 | 23.00% |
| D6_1 | 305 | 5004 | 94 | 23.56% |
| D6_2 | 326 | 5000 | 100 | 23.47% |
| D6_3 | 303 | 4980 | 89 | 22.70% |
| D10_1 | 338 | 4900 | 100 | 22.83% |
| D10_2 | 309 | 4986 | 95 | 23.51% |
| D10_3 | 336 | 4997 | 101 | 23.11% |
| D15_1 | 326 | 5001 | 99 | 23.29% |
| D15_2 | 306 | 5008 | 92 | 23.12% |
| D15_3 | 329 | 4950 | 98 | 22.95% |

*Table S3*. Relative distribution of GO functional classification of the identified surface proteins during chondrogenic differentiation in micromass cultures

| | Day 1 | Day 3 | Day 6 | Day 10 | Day 15 |
|---|---|---|---|---|---|
| Receptors | 28% | 29% | 27% | 28% | 28% |
| Enzymes | 33% | 31% | 32% | 32% | 31% |
| Transporters | 21% | 21% | 19% | 19% | 20% |
| Adhesion proteins | 24% | 23% | 23% | 23% | 23% |
| Miscellaneous | 20% | 24% | 25% | 25% | 26% |
